# Targeted CRISPR Screens Reveal Genes Essential for *Cryptosporidium* Survival in the Host Intestine

**DOI:** 10.1101/2024.11.22.624643

**Authors:** Lucy C Watson, Katarzyna A Sala, Netanya Bernitz, Lotta Baumgärtel, Mitchell A Pallett, N Bishara Marzook, Lorian Cobra Straker, Duo Peng, Lucy Collinson, Adam Sateriale

## Abstract

The *Cryptosporidium* parasite is one of the leading causes of diarrheal morbidity and mortality in children, and adolescent infections are associated with chronic malnutrition. There are no vaccines available for protection and only one drug approved for treatment that has limited efficacy. A major barrier to developing new therapeutics is a lack of foundational knowledge of *Cryptosporidium* biology, including which parasite genes are essential for survival and virulence. Here, we iteratively improve the tools for genetically manipulating *Cryptosporidium* and develop a targeted CRISPR-based screening method to rapidly assess how the loss of individual parasite genes influence survival *in vivo*. Using this method we examine the parasite’s pyrimidine salvage pathway and a set of leading *Cryptosporidium* vaccine candidates. From this latter group we determined the parasite gene known as Cp23 to be essential for survival, which was confirmed through inducible knockout *in vitro* and *in vivo*. Parasites deficient in Cp23 were able to replicate within and emerge from infected epithelial cells, yet unable to initiate gliding motility which is essential for the reinfection of neighbouring cells. The targeted screening method presented here is highly versatile and will enable researchers to more rapidly expand the knowledge base for *Cryptosporidium* infection biology, paving the way for new therapeutics.

## Introduction

Diarrhoeal related infections are a major cause of morbidity and mortality in children around the world (Troeger et al., 2017), (Troeger et al., 2018). Cryptosporidiosis is consistently found to be one of the leading causes of a moderate-to-severe diarrhoeal disease in infants (Kotloff et al., 2013), (Kotloff et al., 2019). Unlike other diarrheal diseases that are attributed with high incidence rates, such as rotavirus or *Shigella*, there are no effective drugs or vaccines for *Cryptosporidium*. Nitazoxanide, the only Food and Drug Administration approved drug for treatment, is not effective in immunocompromised individuals, only partially effective in adults, and not approved for use in children, the patient population that needs intervention the most (Abubakar et al., 2007), (Amadi et al., 2009). One reason for this scarcity of therapeutics is a historic lack of effective systems to study the parasite. While there have been recent improvements in genetic manipulation (Vinayak et al., 2015), drug target identification (Caldwell et al., 2024), (Manjunatha et al., 2024), (Ajiboye et al., 2024), and animal models of infection (Sateriale et al., 2019), our comprehension of *Cryptosporidium* biology remains very basic. Specifically, there is a limited understanding of the *Cryptosporidium* genes that contribute to parasite fitness and survival, genes that would be the most suitable targets for therapeutic intervention.

Reverse genetic approaches have been instrumental in identifying parasite genes that influence survival and convey fitness in other Apicomplexan parasites (Sidik et al., 2016), (Young et al., 2019), (Bushell et al., 2017), (Butterworth et al., 2023), (Smith et al., 2022). *Toxoplasma*, which has historically served as a facile model for Apicomplexa, has benefited from a high transfection efficiency coupled with non-homologous end joining (NHEJ) to be at the forefront of Apicomplexa CRISPR screening (Sidik et al., 2016). Like *Cryptosporidium*, the *Plasmodium* parasite lacks NHEJ pathways, so CRISPR-Cas9 driven double stranded breaks can only be fixed by homologous repair. This lack of NHEJ can be leveraged to implement precise CRISPR screening, yet to date, no such screening method exists in *Cryptosporidium*. Here, we overcome the current technical barriers for *Cryptosporidium* genetic manipulation to develop a reproducible method for pooled *in vivo* CRISPR screens.

*Cryptosporidium* has a highly streamline genome and lacks many basic metabolic pathways. Because of this, it is thought to be heavily reliant on its host cell for nutrients. This is particularly evident in the nucleotide salvage pathway, where *Cryptosporidium* is believed to be capable of scavenging both nucleotides and nucleotide precursors from the host enterocyte (Pawlowic et al., 2019). Despite a compact genome, both purine and pyrimidine salvage pathways show evidence of redundancy. The purine salvage pathway has been well studied in *Cryptosporidium*, revealing that many genes in the pathway can be independently eliminated, resulting in little or no deficit in parasite growth (Pawlowic et al., 2019). Surprisingly, this includes inosine monophosphate dehydrogenase (IMPDH), a gene once considered to be a crucial drug target based on metabolic mapping (Striepen et al., 2004), (Jefferies et al., 2015). The absence of a growth defect in parasites that lack IMPDH suggests that *Cryptosporidium* can not only synthesise purine nucleotides, but also take them up directly from the infected host cell. Less is known about the pyrimidine salvage pathway in *Cryptosporidium*, however, there is still evidence of redundancy. Thymidine kinase (TK) and dihydrofolate reductase-thymidine synthase (DHFR-TS) both convert their respective substrates (deoxythymidine or deoxyuridine monophosphate) to deoxythymidine monophosphate. For this reason, TK and DHFR-TS are both independently non-essential for DNA synthesis and parasite growth (Vinayak et al., 2015), (Pawlowic et al., 2019). Using our *in vivo* pooled CRISPR screen, we discovered that most of the pyrimidine salvage genes significantly contribute to parasite fitness within the intestine.

It is widely accepted that protective immunity can be acquired to *Cryptosporidium*, as incidences of *Cryptosporidium* infections decrease with age (Haque et al., 2009), (Kotloff et al., 2013), (Sow et al., 2016), and experimental infection with attenuated parasites leads to protection in calves and mice (Jenkins et al., 2004), (Sateriale et al., 2019). Considering the success of the vaccination campaign for rotavirus (Yen et al., 2014), (O’Ryan, 2017), a diarrheal pathogen with a similar patient population and pathogenesis, a *Cryptosporidium* vaccine is predicted to greatly reduce morbidity and mortality in children. Numerous protein antigens have been associated with protection in humans and several prominent surface proteins have been suggested as vaccine candidates (Gilchrist et al., 2023), (Manque Patricio et al., 2011), (Askari et al., 2016). Yet, if and how many of these proteins contribute to parasite fitness is unclear. Here, we assess the relative fitness contributions during infection for a panel of proposed *Cryptosporidium* vaccine candidates (collected from the literature). As Cp23 (also known as the *Cryptosporidium* immunodominant antigen 23) is one of the leading vaccine candidates, and strongly correlated with protection in humans (Gilchrist et al., 2023), we investigated how this protein specifically contributes to parasite virulence and fitness.

## Results

### Iterative improvements to improve *Cryptosporidium* genetic manipulation

To develop CRISPR screening, we first sought to improve *Cryptosporidium* transfection efficiency. To transfect *Cryptosporidium*, dormant oocysts are treated with bile salts and warmed to body temperature to simulate the conditions of a human intestine. Under these conditions, the environmentally hardy oocysts ‘excyst’, releasing four motile and transfectable sporozoites that readily invade intestinal epithelial cells. To optimise transfection, we transfected *Cryptosporidium parvum* parasites with a vector that contained both luminescent (NanoLuciferase) and fluorescent (mCherry) reporters under constitutive expression and allowed the transfected parasites to infect an intestinal cell (HCT8) monolayer. Numerous electroporation programs were trialled using transient expression, with FL115 demonstrating the highest luminescence **(Sup. 1a).** Furthermore, various combinations of bile salts and incubation media were tested for their effect on parasite transfection. Notably, incubation in sodium taurocholate led to higher transfection efficiency compared to the deoxy form, sodium taurodeoxycholate **(Sup. 1b)**. Combined, these adjustments resulted in more than a 50-fold improvement in transfection efficiency **(Sup. 1c & d)**.

As *Cryptosporidium* parasites lack NHEJ, CRISPR driven injury of the genome is required to drive homologous recombination for genetic manipulation. Although this represents a significant barrier to high-throughput screening, the parasite’s lack of NHEJ very likely contributes to its uniquely high level of genetic editing specificity. Indeed, an ‘oK-target’ genomic insertion has yet to be reported by *Cryptosporidium* researchers. Further, the parasite predominantly exists in a haploid state during its life cycle and has a minimal requirement of homology for efficient editing. To determine the smallest reliable allowance for efficient repair, we tested the efficiency of editing using differing length homology arms to integrate a nanoluciferase gene into the dispensable TK locus. A positive correlation between the length of the homology arms and the efficiency of repair was observed, with 50bp (the longest length tested) being the most efficient, while no integration was observed in the absence of CRISPR driven injury of TK (**Sup. 1e**).

To allow for pooled genetic knockout screens, we reasoned that a one-plasmid approach would be required, where a single plasmid delivers the Cas9-expression vector, a target specific gRNA, and a segment of DNA to be inserted at the target genomic locus (repair DNA). The repair DNA contains homologous flanks surrounding an expression cassette with the selectable marker and a genetic barcode for identification. Further, to enable high-throughput creation of targeted libraries, we developed a two-step Golden Gate assembly method, where first a 300bp segment containing all gene-specific (unique) DNA is integrated into the Cas9-expression vector. In the second step of the assembly, a non-unique expression cassette is inserted that will integrate into the genome, enabling the use of virtually any combination of selection markers or reporters **(Fig. 1a)**. To fit all unique DNA into a 300bp oligonucleotide, we used a 50bp gRNA which simultaneously serves as one of the homology sites for genomic integration. To test the feasibility of this approach, we generated a vector targeting TK, a gene that is known to be dispensable for parasite growth. The TK targeting vector was transfected into *Cryptosporidium parvum (C. parvum)* sporozoites that were used to infect genetically immunocompromised (interferon gamma deficient *Ifnγ^-/-^*) mice and luminescence was measured in mouse faecal material 7 days post transfection, comparable to what is observed when using a separate DNA repair template and a Cas9-expression vector **(Fig. 1b)**. As we redefined the method to generate *Cryptosporidium* transgenics, we confirmed the approach’s specificity via whole genome sequencing, and as expected, the only genome alteration observed was at the targeted site within the coding region of the TK gene **(Fig. 1c)**.

**Figure 1.**
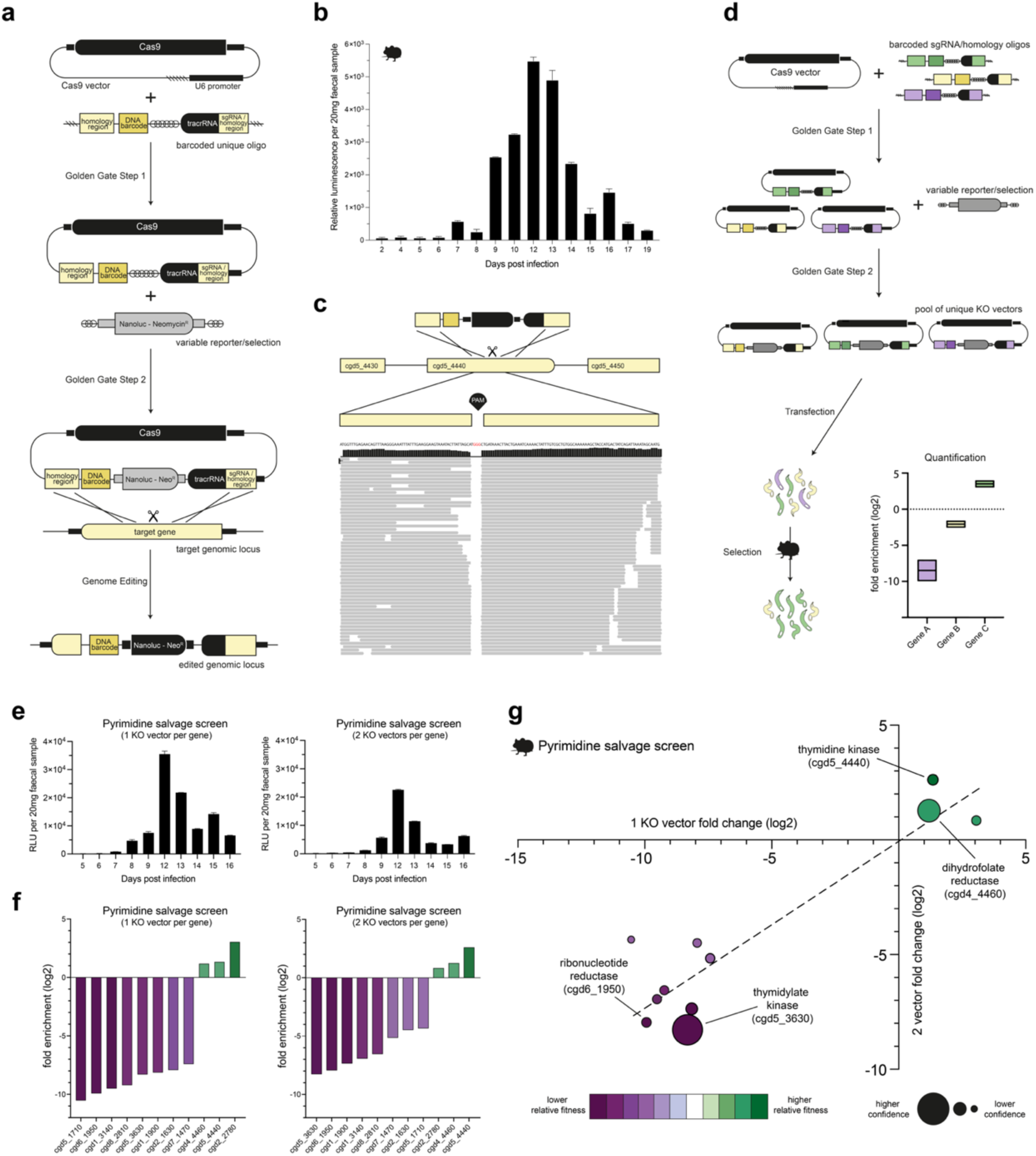
*In vivo* CRISPR Screen of the *Cryptosporidium* Pyrimidine Salvage Pathway. **a.** Schematic illustrates how a knockout vector is generated. The first Golden Gate reaction occurs between a Cas9 expression plasmid and a 300bp unique segment (containing the two 50bp homology arms (one of which serves dual function as the gRNA), a unique DNA barcode, BsmBI restriction enzyme sites and the tracrRNA). The second Golden Gate reaction, using the BsmBI restriction enzyme sites, inserts a variable selection/reporter cassette to generate a complete knockout vector. The knockout vector then contains all the machinery to disrupt a gene of interest by inserting a variable selection cassette and barcode at the genomic locus. **b.** A knockout vector targeting thymidine kinase (cgd5_4440) was generated and transfected into *C. parvum* sporozoites that were used to infect Ifnγ^-/-^ mice under paromomycin selection. Faecal samples were collected and the luminescence in faecal material was monitored. Data shows the mean of a biological replicate ± SEM of the 2 technical replicates, n = 4 mice. **c.** Alignment of reads from whole genome sequencing of the thymidine kinase knockout strain to the *C. parvum* IOWAII genome at the site of insertion. Note the complete lack of alignment to the PAM site, which is removed by the homologous repair event. **d.** Overview of CRISPR screening method. Following construction of KO vectors (detailed in 1a), sporozoites are transfected with gene specific KO vectors and used to infect mice. Specific barcodes are then amplified via high fidelity PCR and used to calculate fold enrichment (log2[%barcode(output) / %barcodes(input)]) which measures the relative fitness contribution of each gene. **e.** Mouse faecal material was collected, and luminescence was monitored in the pooled sample from the pyrimidine salvage pathway CRISPR screens at 1 and 2 KO vectors per gene. Data shown is the mean of the biological replicate ± SEM of the 2 technical replicates, n = 4 mice per screen. **f.** Rank ordered fold enrichment scores from the 1 and 2 KO vectors per gene pyrimidine salvage CRISPR screens. The colour indicates the relative fitness contribution, with dark purple showing high fitness conferring and dark green showing low fitness conferring. **g.** Comparison of the fold enrichment scores between the 1 and 2 KO vectors per gene pyrimidine salvage CRISPR screens. Confidence refers to the inverse of the 95% confidence interval when comparing the log2 fold change scores from each screen (see methods).

### CRISPR KO screen of the pyrimidine salvage pathway

To assess the feasibility and reproducibility of a recombination-based *in vivo* screen, we designed a pilot screen targeting 11 genes in the parasite’s pyrimidine salvage pathway (**Table. 1**). We ran parallel screens employing either 1 or 2 targeting vectors per gene, hence 11 or 22 gRNAs, respectively. Targeted KO vectors were transfected into *C. parvum* sporozoites, which were propagated in *Ifnγ^-/-^* mice under paromomycin selection **(Fig. 1d)**. Luminescence from the integrated nanoluciferase reporter was detected in mouse faecal material 8 days post transfection, in both the 1 and 2 KO vector per gene pyrimidine salvage screens **(Fig. 1e)**. From the output, faecal material from the infected mice, parasites were purified, their DNA extracted, and the barcodes amplified via high fidelity PCR. From the input, transfected parasites used to infect the mice, DNA was extracted and the barcodes amplified via high fidelity PCR. Both the output and input barcodes were sequenced and used to calculate fold enrichment scores for each gene. Fold enrichment scores serve as a measure of relative fitness, genes whose barcodes show a negative enrichment are presumed to be important to parasite survival and therefore highly fitness conferring **(Fig. 1f)**. Importantly, fold enrichment scores between the 1 and 2 KO vector screens were strongly correlated, achieving an R^2^ of 0.842 **(Fig. 1g)**. Of the 11 genes included in the pyrimidine salvage screen, only 3 were low fitness conferring: dihydrofolate reductase – thymidylate synthase (DHFR-TS) (cgd4_4460), thymidine kinase (TK) (cgd5_4440) and dCMP deaminase (cgd2_2780). As mentioned previously, TK and DHFR-TS play redundant roles in synthesis of *Cryptosporidium* dTMP, making them independently non-essential for DNA synthesis and parasite growth. Thus, the results of our pilot screen effectively mirror those of previous experiments attempting individual knockouts of these genes *in vivo* (Vinayak et al., 2015), (Pawlowic et al., 2019).

**Table 1.**
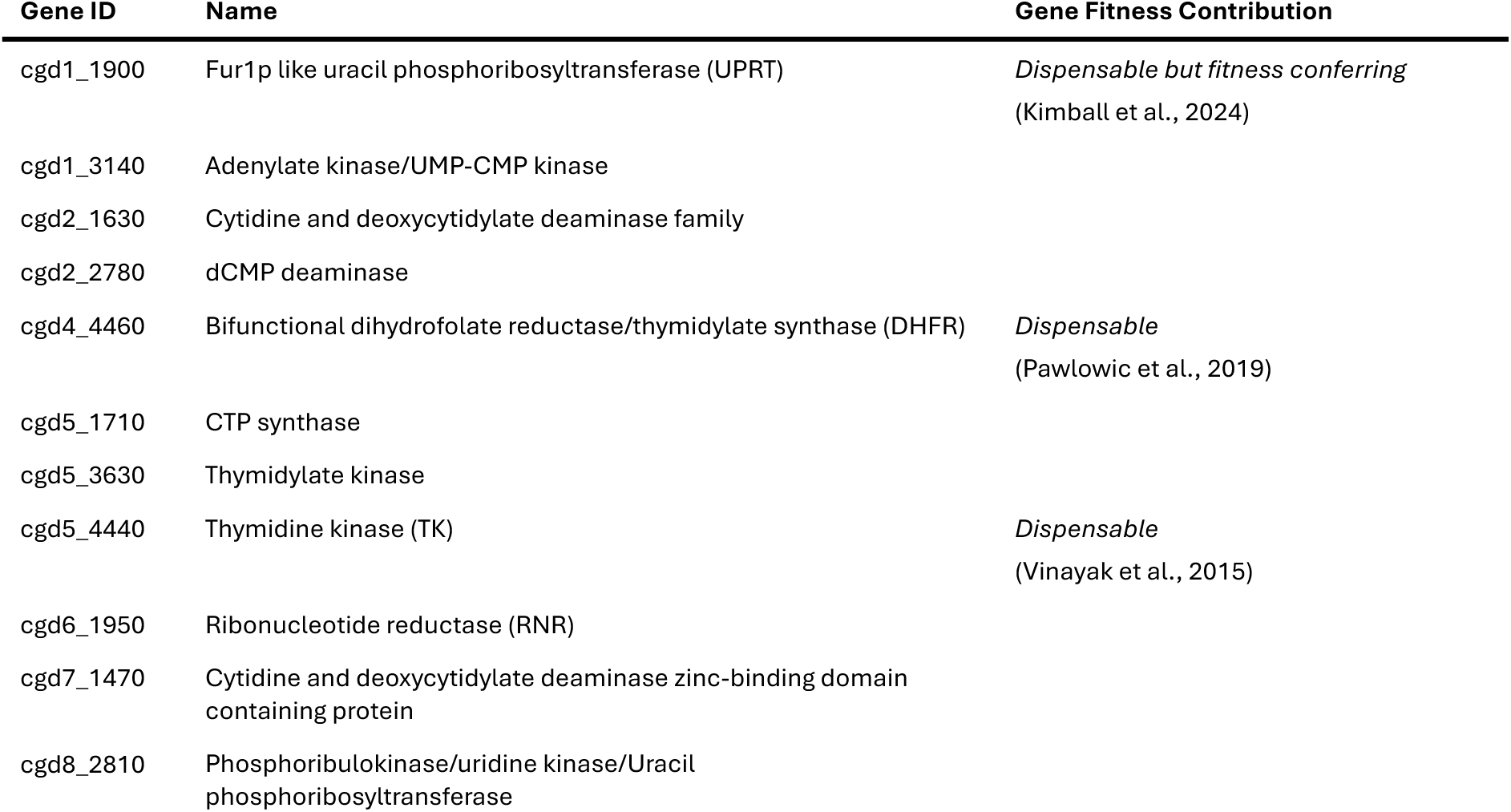
Genes in the Pyrimidine Salvage CRISPR Screen.

| Gene ID | Name | Gene Fitness Contribution |
| --- | --- | --- |
| cgd1_1900 | Fur1p like uracil phosphoribosyltransferase (UPRT) | *Dispensable but fitness conferring*<br>(Kimball et al., 2024) |
| cgd1_3140 | Adenylate kinase/UMP-CMP kinase |  |
| cgd2_1630 | Cytidine and deoxycytidylate deaminase family |  |
| cgd2_2780 | dCMP deaminase |  |
| cgd4_4460 | Bifunctional dihydrofolate reductase/thymidylate synthase (DHFR) | *Dispensable*<br>(Pawlowic et al., 2019) |
| cgd5_1710 | CTP synthase |  |
| cgd5_3630 | Thymidylate kinase |  |
| cgd5_4440 | Thymidine kinase (TK) | *Dispensable*<br>(Vinayak et al., 2015) |
| cgd6_1950 | Ribonucleotide reductase (RNR) |  |
| cgd7_1470 | Cytidine and deoxycytidylate deaminase zinc-binding domain containing protein |  |
| cgd8_2810 | Phosphoribulokinase/uridine kinase/Uracil phosphoribosyltransferase |  |

### Refining diCRE-mediated genetic editing in *Cryptosporidium*

To validate our results from the pyrimidine salvage screen, we sought to use an inducible Cre-recombinase system to remove genes at their genetic locus. This method has been previously adapted for use in *C. parvum*, employing the conditional split recombinase (diCRE), which dimerises upon rapamycin addition and excises DNA between loxP recombination sites, one of which is embedded within an artificial intron introduced into the target gene’s coding sequence (Tandel et al., 2023), (Shaw et al., 2024). This system demonstrated high efficiency, but low-to-moderate activity in the absence of the rapamycin induction (i.e. leakiness). To refine this system, we tested a validated intron from the male gamete fusion factor HAP2 (cgd8_2220). Insertion of this intron within the nanoluciferase reporter with and without the loxP recombination site did not affect the measured luminescence during *in vitro* infection **(Sup. 2a)**. Using this validated intron/loxP combination, we generated a stable *C. parvum* diCRE parasite line targeting TK with the diCRE subunits, FRB-Cre60 and FKBP-Cre59, expressed under the same promoter, separated via a self-cleaving T2A skip peptide (TK-T2A-diCRE) **(Sup. 2b)**. However, when investigating the excision dynamics of this *C. parvum* TK-T2A-diCRE line, by infecting HCT8 monolayers in the presence or absence of rapamycin, we noted a high level of excision in the absence of rapamycin, similar to what has been reported previously **(Sup. 2c)**. We reasoned that the T2A skip peptide may not be functioning properly and causing background induction in *Cryptosporidium*. Consequently, we generated a stable transgenic parasite line targeting TK where the diCRE segments were under independent aldolase and tubulin promoters (TK-diCRE) **(Sup. 2d)**. When investigating the excision dynamics of this TK-diCRE line, we observed complete excision of the loxP-flanked TK segment at 24 hours post rapamycin induction, and no measurable excision in the non-induced controls **(Sup. 2e).** Using these TK-diCRE parasites **(Sup. 3a)**, we performed a time course to measure the dynamics of knockout at both the DNA and protein level (via a C-terminal HA tag). DNA excision had not occurred by 8 hours post rapamycin treatment but was complete by 24 hours, and no excision was detected in the non-induced control **(Sup. 3b)**. Likewise, protein levels started to decrease by 12 hours post rapamycin treatment and were approximately 95% reduced compared to the non-induced controls by 24 hours. In contrast, the protein levels remained high throughout the time course in the non-induced controls **(Sup. 3c & d)**.

### Ribonucleotide reductase is required for parasite DNA replication and survival

Ribonucleotide reductases (RNRs) catalyse the conversion of nucleotides to deoxynucleotides, an essential step for DNA synthesis in all organisms. In our pyrimidine salvage screen, RNR showed the lowest fold enrichment score, indicating a high effect on parasite fitness. Our first attempt to create RNR-diCRE parasites was unsuccessful, likely due to the addition of a C-terminal HA epitope tag. It has been suggested that the C-terminus of RNR is required for its function in other organisms and this may hold true for *Cryptosporidium* (Cohen et al., 1986), (Dutia et al., 1986). The second attempt to generate RNR-diCRE parasites, without the C-terminal HA epitope tag, was successful **(Sup. 3e)**. *In vitro*, rapamycin induced DNA excision of both TK- and RNR-diCRE parasites appears to follow similar dynamics, although we note some partial excision in the RNR-diCRE parasites at later timepoints (**Fig. 2a & b**). Further, functional deletion of both TK and RNR was confirmed using 5-ethynyl2’-deoxyuridine (EdU), a thymidine analogue. Thymidine kinase is required for EdU phosphorylation and incorporation, and it has been shown before that loss of the TK gene in *Cryptosporidium* leads to a failure to incorporate EdU (Vinayak et al., 2015). In contrast, RNR deletion will not block EdU phosphorylation, but rather incorporation, as DNA synthesis cannot occur without other deoxynucleotides. We observed a complete lack of EdU incorporation in rapamycin treated TK and RNR-diCRE parasites, while EdU incorporation was observed in the non-induced and wildtype controls **(Sup. 3f)**. *In vitro*, deletion of TK led to no growth defect, whereas deletion of RNR strongly attenuated parasite growth by 24 hours post infection **(Fig. 2c)**. Similarly, rapamycin treatment of TK-diCRE infected mice lead to no growth defect, but rapamycin treatment of RNR-diCRE infected mice completely inhibited parasite growth **(Fig. 2d)**, demonstrating that RNR is indeed essential for parasite survival, further underlining the reliability of the *in vivo* CRISPR screening method.

**Figure 2.**
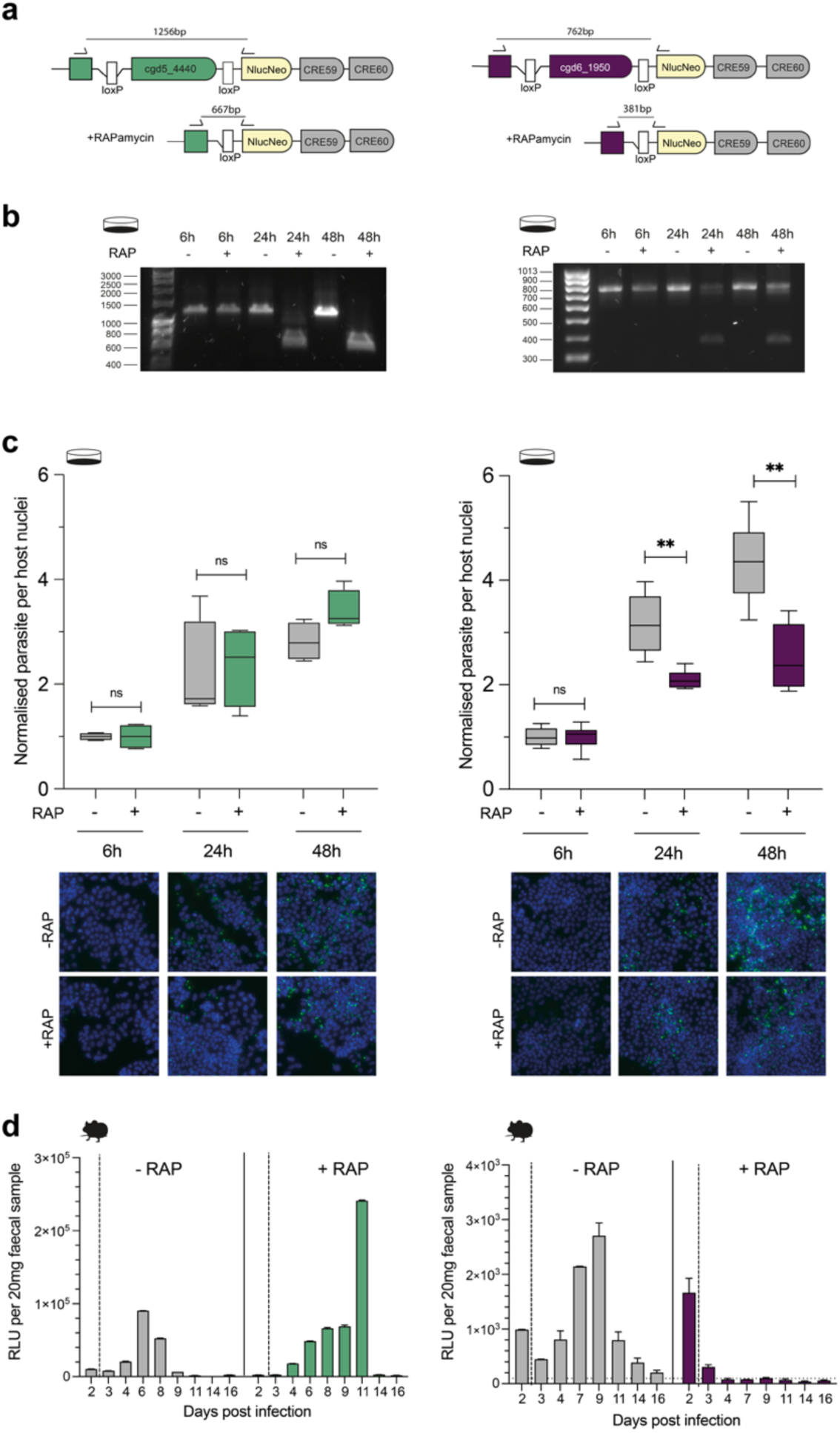
Validating Screen Results with diCRE Mediated Excision. **a-b.** TK-diCRE (shown in green) and ribonucleotide reductase (RNR) -diCRE (shown in purple) parasites were generated and used to infect an HCT8 monolayer in the presence or absence of rapamycin. Genomic DNA was extracted at 6, 24 and 48 hours and diagnostic PCRs confirmed the level of excision at the given time points. **c.** HCT8 monolayers infected with TK-diCRE and RNR-diCRE parasites in the presence or absence of rapamycin. At 6, 24 and 48 hours the monolayers were fixed, stained and the number of parasites per host nuclei was quantified. Representative images are shown, with nuclei in blue (Hoechst), and parasites in green (Vicia villosa lectin). Data shown is representative of 2 biological replicates ± sd of 4 technical replicates. Significance is determined using a two-tailed unpaired t-test, ns = not significant, ** = p ≤ 0.01. **d.** TK-diCRE or RNR-diCRE parasites were used to infect Ifnγ^-/-^ mice, which were either treated with rapamycin or vehicle (DMSO) in their drinking water starting at day 2 post infection. Data shown is a biological replicate ± SEM of the 2 technical replicates, n = 2 mice per condition.

### CRISPR KO screen of *Cryptosporidium* vaccine candidates

Both the 1 and 2 KO vector per gene pyrimidine salvage screens were reproducible, but to circumvent potential low efficiency gRNAs, we chose to screen the leading *Cryptosporidium* vaccine candidates with 2 KO vectors per gene. In total, 11 vaccine candidates (22 KO vectors) were chosen due to their reported surface location and implication of immunogenicity and/or immune protection in the literature **(Table. 2)**. Luminescence from the nanoluciferase reporter was detected in mouse faecal material around 7 days post transfection in both screens **(Fig. 3a)**. As before, parasites were purified, DNA extracted, and barcodes were amplified via high fidelity PCR. The sequenced barcode counts were used to calculate fold enrichment scores for each gene **(Fig. 3b)**, and importantly these scores were comparable, achieving an R^2^ of 0.611 **(Fig. 3c)**. Of the 11 genes in the vaccine candidate screen, only 2 were low fitness conferring: cgd6_1660 (thrombospondin repeat protein 11 (TSP11)) and cgd6_32 (apical glycoprotein 1 (AGP1)). The remaining 9 genes appeared to confer some level of fitness. Akey et al., recently demonstrated that AGP1 was dispensable for parasite survival, while apical glycoprotein 2 (AGP2) (cgd7_4330) was likely essential, both phenotypes that our pooled *in vivo* CRISPR screen recapitulated (Akey et al., 2023).

**Figure 3.**
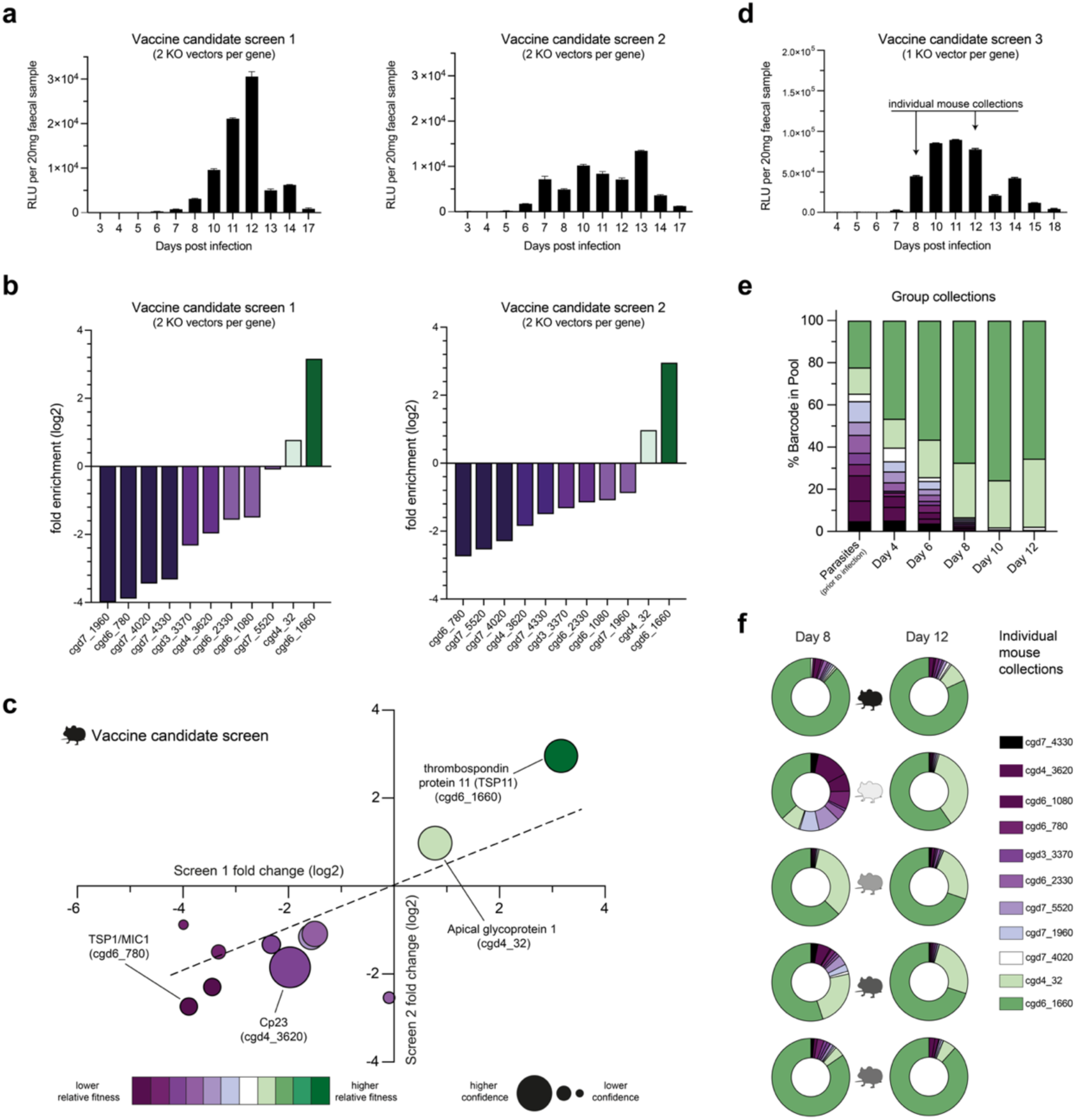
*In vivo* CRISPR Screen of *Cryptosporidium* Vaccine Candidates. **a.** Mouse faecal material was collected and luminescence was monitored during infection. Each replicate was conducted with 2 KO vectors per gene. Data shown is the mean of the biological replicate ± SEM of the 2 technical replicates, n = 5 Ifnγ^-/-^ mice for repeat 1 and n = 3 Ifnγ^-/-^ mice for repeat 2. **b.** Rank ordered fold enrichment scores from the vaccine candidate CRISPR screens. The colour indicates the relative fitness contribution of a gene, with dark purple being high fitness conferring and dark green being low fitness conferring. **c.** Comparison of the fold enrichment scores between the replicate CRISPR screens for each gene. Confidence refers to the inverse of the 95% confidence interval when comparing the log2 fold change scores from each screen (see methods). **d.** Mouse faecal material was collected and luminescence was monitored during infection. Data shows a biological replicate ± SEM of the 2 technical replicates, n = 5 Ifnγ^-/-^ mice. **e-f.** Barcodes from KO parasites could be easily monitored over time (**e**) and within individual mice (**f**).

**Table 2.** Genes in the Vaccine Candidate CRISPR Screen.

| Gene ID | Name | Gene Fitness Contributions |
| --- | --- | --- |
| cgd3_3370 | Uncharacterized protein |  |
| cgd4_32 | Apical glycoprotein 1 (AGP1) | *Dispensable*<br>(Akey et al., 2023) |
| cgd4_3620 | Immunodominant antigen 23393226 (Cp23) |  |
| cgd6_1080 | Glycoprotein GP40 (GP60) | *Dispensable, but fitness conferring*<br>(Li et al., 2024) |
| cgd6_1660 | Uncharacterized protein with Thrombospondin type-1 (TSP1) repeat (TSP11) |  |
| cgd6_2330 | Uncharacterized protein |  |
| cgd6_780 | Thrombospondin type-1 (TSP1) repeat/EGF-like domain containing protein (TSP8/MIC1) |  |
| cgd7_1960 | WD40/YVTN repeat-like+signal peptide-containing protein |  |
| cgd7_4020 | Cryptosporidial mucin (GP900) |  |
| cgd7_4330 | Apical glycoprotein 2 (AGP2) | *Likely Essential*<br>(Akey et al., 2023) |
| cgd7_5520 | Hemogen |  |

To gain more insight into growth and competition of pooled KO parasites, we conducted an additional 1 KO vector per gene vaccine candidate screen, where instead of amplifying DNA barcodes from parasites purified around the peak of infection, we amplified barcodes directly from individual faecal collections **(Fig. 3d)**. This alternative method offered greater temporal and spatial resolution of infection where fold enrichment scores could be easily and non-invasively be monitored throughout the experiment from either pooled collections or individual mice **(Fig. 3e & f)**. Individual mice demonstrated considerable heterogeneity at day 8 of infection, which decreased over time and became more uniform by day 12. Again, this screen found that AGP1 and TSP11 were low fitness conferring relative to the other genes within the cohort.

### Immunodominant antigen 23 is required for host cell invasion

Immunodominant antigen 23 (Cp23) (cgd4_3620) is one of the leading *Cryptosporidium* vaccine candidates, having been originally discovered in 1986 (Ungar & Nash, 1986). Despite nearly 40 years of research, there is very little known about Cp23’s role and function during *Cryptosporidium* infection. In our vaccine candidate screen, Cp23 displayed a low fold enrichment score, indicating a moderate-to-high effect on parasite fitness. To investigate the function of Cp23, we generated Cp23-diCRE parasites **(Sup. 4a)**. *In vitro*, the rapamycin induced DNA excision had started by 6 hours and was complete by 24 hours, with no excision observed in the uninduced controls **(Fig. 4a & b)**. Loss of Cp23 was confirmed at the protein level using a commercially available monoclonal antibody. *In vitro*, rapamycin treatment caused a significant decrease in parasite growth after 24 hours **(Fig. 4c)** and rapamycin treatment of Cp23-diCRE infected mice completely blocked parasite growth **(Fig. 4d)**, demonstrating that Cp23 is indeed essential for parasite survival.

**Figure 4.**
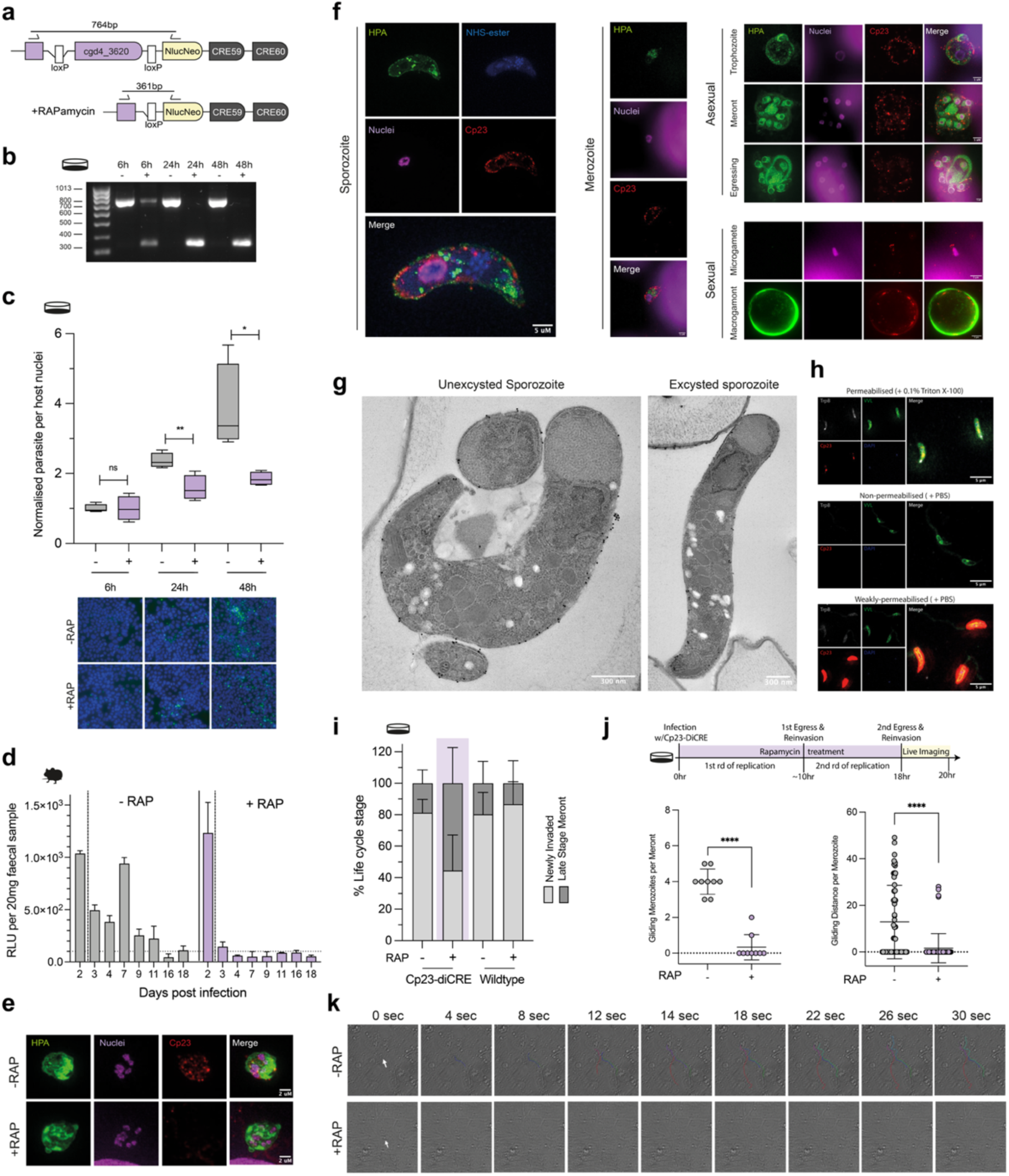
Immunodominant Antigen 23 is Essential and Required for Reinvasion of Host Cells. **a-b.** HCT8 cell monolayers infected with Cp23-diCRE in the presence or absence of rapamycin. At 6, 24 and 48 hours, gDNA was extracted and diagnostic PCRs confirmed the level of excision at the given timepoint. **c.** HCT8 cell monolayers infected with Cp23-diCRE in the presence or absence of rapamycin. At 6, 24 and 48 hours, the monolayer was fixed and stained and the parasite per host nuclei was quantified. Representative images are shown, with nuclei in blue (Hoechst), and parasites in green (Vicia villosa lectin). Data shown is representative of two biological replicates ± sd of the 4 technical replicates. Significance is determined using a two-tailed unpaired t-test, ns = not significant, * = p ≤ 0.05, ** = p ≤ 0.01**. d.** Ifnγ^-/-^ mice were infected with 50,000 Cp23-diCRE parasites and treated with rapamycin or DMSO control in their drinking water at 2 days post infection. Data shown is the mean of a biological replicate ± SEM of the 2 technical replicates, n = 2 mice per condition. **e.** Super resolution microscopy at 24 hours post infection *in vitro* with Cp23-diCRE parasites in the presence or absence of rapamycin; green (helix pomatia agglutinin (HPA)), parasite; magenta, (sytox), nuclei; red (Cp23 Ab), Cp23. Scale = 2μm. **f.** Expansion microscopy of the *C. parvum* asexual stages: sporozoite, merozoite, trophozoite and meront, and the sexual stages: macrogamont (female) and microgamete (male); green (helix pomatia agglutinin (HPA)); blue (N-hydroxysuccinimide (NHS ester); magenta (sytox); red (αCp23). Scale bar = 5μm. Median expansion factor of 4.5. **g.** Transmission electron microscopy of an excysted and unexcysted *C. parvum* sporozoite coupled with immunogold labelling with Cp23 antibody. Scale = 300nm**. h.** Permeabilisation assay of *C. parvum* sporozoites. The permeabilised condition used 0.1% Triton X-100 and the non-permeabilised condition used PBS. Grey (tryptophan synthase (αTrpB); green (Vicia villosa lectin); red (αCp23); blue (Hoechst). All images were taken using the same settings and exposure. Scale bar = 5μm. **i.** HCT8 cell monolayers were infected with either Cp23-diCRE or wildtype parasites in the presence or absence of rapamycin, and at 22 hours post infection life stages were quantified. Data shown is 2 biological replicates, n = 438 Cp23-diCRE - RAP, n = 173 Cp23-diCRE + RAP, n = 213 wildtype - RAP, n = 203 wildtype + RAP. Significance is determined using a two-tailed unpaired t-test, ns = not significant, **** = p ≤ 0.0001. **j-k.** Live imaging of the reinvasion event was carried out at 18 hours post infection of an HCT8 monolayer with Cp23-diCRE parasites in the presence or absence of rapamycin. Movement of merozoites was tracked using Fiji software. Representative images are shown in **k.** with videos in supplementary data. All live microscopy data shown is from 2 biological replicates. Significance is determined using a two-tailed unpaired t-test, **** = p ≤ 0.0001.

Cp23 has been reported to be on the surface of the sporozoite (Mead et al., 1988), in the sporozoite trails (Arrowood et al., 1991), and is predicted to localise to the micronemes (Guérin et al., 2023). To localise Cp23 at high resolution, we used a commercial antibody that we genetically validated with our Cp23-diCRE parasites **(Fig. 4e)**, coupled with expansion and super-resolution microscopy. This revealed Cp23 was expressed at the parasite’s pellicle throughout the life cycle **(Fig. 4f)**. To resolve the location further, we carried out transmission electron microscopy (TEM) with the immunogold labelled Cp23 antibody **(Fig. 4g)**. In excysted and unexcysted sporozoites, Cp23 again appeared to primarily localise to the parasite pellicle, either at the plasma membrane or inner membrane complex (IMC). The immunogold-TEM also revealed that even in unexcysted sporozoites, Cp23 was not present in the parasite’s micronemes. Further, we found no evidence that Cp23 was present in sporozoite trails **(Sup. 4b)**. When permeabilised with Triton X-100, sporozoites demonstrated faint staining with the Cp23 antibody **(Fig. 4h)**. Without permeabilisation, *C. parvum* sporozoites, surprisingly, demonstrated either a complete lack of Cp23 signal or a heighted intensity of signal throughout the parasite **(Fig. 4h)**. In these high intensity parasites, we can detect the internal control antibody (CpTrpB - *Cryptosporidium* tryptophan synthase beta) at low levels. This suggests that 1) high intensity Cp23 parasites are weakly permeabilised, 2) permeabilisation with Triton X-100 affects Cp23 localisation, likely by disrupting its membrane association, and 3) Cp23 is likely not exposed on the surface of the sporozoite.

Within our synchronised *in vitro* model of infection, *C. parvum* parasites egress from their infected cell and re-invade nearby epithelial cells around 18 hours post infection, starting another round of asexual reproduction. By 22 hours, this reinvasion event is mostly complete. To determine the biological function of Cp23, we examined our Cp23-diCRE parasites in the context of this reinfection event, when DNA excision and depletion of Cp23 has started. At 22 hours, Cp23-diCRE parasites without rapamycin treatment were mostly newly invaded life stages (1n or 2n = 81% of parasites observed, n = 356/438). When Cp23 was ablated by rapamycin, fewer parasites observed were newly invaded life stages (1n or 2n = 45%, n = 77/173) **(Fig. 4i & Sup. 4c & e)**. In wildtype controls, both in the presence and absence of rapamycin, again, most of the life stages observed were newly invaded (1n or 2n = 83%, n = 176/203 and 171/213 respectively) **(Fig. 4i & Sup. 4d & e)**. We hypothesised that the reduced percentage of newly invaded parasites when

Cp23 was ablated might be due to a defect in reinvasion. To examine this more closely, live microscopy was carried out in the presence and absence of rapamycin **(Fig. 4j & k, supplementary videos S1-4)**. When Cp23 was ablated, merozoites could egress, but were then unable to move to another cell to initiate reinvasion. Apicomplexan parasites, such as *Cryptosporidium*, are known to use a method of locomotion called gliding motility, where parasites secrete proteins that are then bound by their own surface receptors, allowing for forward propulsion using an actomyosin-based complex (referred to as the glideosome) (Keeley & Soldati, 2004). Loss of Cp23 in *Cryptosporidium* appears to specifically block this gliding motility that is essential for reinfection.

## Discussion

Reverse genetic screens play an important role in molecular biology and have transformed the field for many pathogens. Here, we overcome the current technical barriers to develop a pooled *in vivo* CRISPR KO method that allows for reverse genetic screening in *Cryptosporidium.* As *Cryptosporidium* lacks the molecular machinery for NHEJ, our method had to employ homologous recombination to create stable transgenics. We leveraged two important features of *Cryptosporidium* genetics: 1) the parasite spends nearly all of its life cycle in a haploid form, thus requiring only one recombination event, and 2) there is a minimal length requirement for homologous recombination, in fact 30bp of DNA flanking the Cas9 cut site appears to be sufficient in our experiments. Our two-step cloning approach for generating these targeted KO vectors is designed to be versatile, allowing for an unlimited range of selection markers and reporters that can be inserted into the genome, which will allow researchers to develop new and innovative screens.

Within an infected host, *Cryptosporidium* undergoes multiple rounds of asexual reproduction before differentiating into sexual forms (male and female) that unite to allow for genetic recombination. During the sexual cycle, transgenic parasites in these CRISPR screens can mate and this has the potential to influence results. This risk is partially mitigated by two factors: 1) the parasite must complete at least three rounds of asexual replication prior to sexual differentiation and potential recombination. This gives ample time for any genes that are detrimental to parasite fitness to exert their effect; 2) *Cryptosporidium* has a relatively high rate of recombination, which should allow for genes, even those in close proximity, to segregate in a nearly random fashion (Kimball et al., 2024). In both the pyrimidine salvage and vaccine candidate screens, we identified high and low fitness conferring genes in close proximity to each other **(Sup. 5a)**. Despite these mitigating factors, there is the potential for sublethal gene knockouts to exert synergistic or antagonistic effects and this possibility must be considered during the design process and while analysing and interpreting results.

Another important factor to consider is the amplifying effect of relative fitness screens within an *in vivo* model of infection. Genes that have a mild effect on parasite survival and fitness can be outcompeted within a larger pool. This appears to be the case for uracil phosphoribosyltransferase (UPRT, cgd1_1900), which catalyses the conversion of uracil and phosphoribosylpyrophosphate to uridine monophosphate. UPRT has recently been shown to be non-essential to parasite survival, yet the authors noted that their UPRT KO *Cryptosporidium* parasites appeared to have a growth defect (Kimball et al., 2024). In our *in vivo* screens, UPRT had a negative fold enrichment, indicating that UPRT contributes highly to parasite fitness and this likely reflects the observed growth defect. In contrast to UPRT, dCMP deaminase (cgd2_2780) had little effect on parasite fitness. dCMP deaminase catalyses the conversion of deoxycytidine-monophosphate (dCMP) to deoxyuridine-monophosphate (dUMP), which is then transformed into deoxythymidine-monophosphate (dTMP) by DHFR-TS. Thymidine kinase can also produce dTMP, likely explaining why the loss of dCMP deaminase had little effect on parasite fitness.

Most of the genes in the vaccine candidate CRISPR screen were identified as fitness conferring, with the exception of apical glycoprotein 1 (AGP1) and thrombospondin repeat protein 11 (TSP11). Akey et al. previous showed AGP1 was dispensable for parasite survival and AGP2 was likely essential, both phenotypes that were recapitulated within our screens (Akey et al., 2023). Recently, it has been suggested that there may be redundancy in the *Cryptosporidium* thrombospondin protein family (TSPs), as there are 12 members, many with similar domain structures (John et al., 2023). Although the *Cryptosporidium* TSPs currently have an unknown function, orthologous Apicomplexan TSP genes play a role in adhesion and motility. We included two TSPs in the vaccine candidate screen: 1) TSP8 (cgd6_780, also known as MIC1) and 2) TSP11 (cgd6_1660). TSP8/MIC1 is a known micronemal protein (Putignani et al., 2008) and was identified as high fitness conferring. In contrast, TSP11 was dispensable, demonstrating that there is at least some redundancy within the *Cryptosporidium* TSP protein family. We also confirmed GP40 (cgd6_1080, also known as GP60) and GP900 (cgd7_4020), antigens that are associated with protection from reinfection in humans, influence parasite fitness *in vivo* (Gilchrist et al., 2023).

Cp23 is one of the leading cryptosporidiosis vaccine candidates, and here we demonstrated that it is highly fitness conferring, both *in vitro* and *in vivo*. Cp23 was first identified in 1986 (Ungar & Nash, 1986), and recent studies have revealed Cp23 recognising IgA and IgG are correlated with protection against *Cryptosporidium* infection in humans (Gilchrist et al., 2023). Until now, the function and precise localisation of Cp23 has been unclear. Here we validate a commercial Cp23 monoclonal antibody using diCRE mediated excision of the target gene, then use a combination of ultrastructure expansion, super-resolution, and transmission electron microscopy, to describe the localisation of Cp23 throughout the parasite life cycle at high resolution. Although our localisation agrees with previously published studies that detects Cp23 at the pellicle (Enriquez & Riggs, 1998), our data suggests that Cp23 may not be exposed at the surface of the sporozoite. Cp23 has no detectable signal peptide, transmembrane domain or GPI-anchor, yet was recently demonstrated to be both myristoylated and palmitoylated (Haserick et al., 2017). Myristylation is a non-reversible lipid modifications that is commonly found in proteins that are anchored to internal membranes in other Apicomplexan parasites (Schlott et al., 2018), (Broncel et al., 2020). There are, however, some exceptions to this rule. TgMIC7 is a micronemal protein in *Toxoplasma* that is myristoylated prior to trafficking to the parasite surface (Broncel et al., 2020). Yet, TgMIC7 contains a transmembrane domain that likely facilitates entry into the secretory pathway, whereas Cp23 does not. While we cannot rule out trafficking of Cp23 to the outer membrane of the parasite, our data supports an internal localisation.

Despite this internal localisation, it is clear Cp23 IgA and IgG have both been correlated with protection in the clinic (Gilchrist et al., 2023). One interpretation of this correlation is that antibodies recognising Cp23 arise from severe *Cryptosporidium* infection(s) where cell-based adaptive immunity against the parasite has developed. Controlled experiments using a natural murine model of *Cryptosporidium* suggest that antibody-based immunity is dispensable for resolution of infection (Sateriale et al., 2019). Further, we found that the genetically validated Cp23 antibody used in this study was unable to block parasite attachment or invasion *in vitro* **(Sup. 6a - b)**. However, it has been reported that antibodies to Cp23 may have a protective effect *in vivo*, thus we cannot definitively rule out the possibility of Cp23-directed protection, whether through antibody or cell-based immunity (Enriquez & Riggs, 1998). Certainly, more investigation is warranted given we now know Cp23’s essential role in parasite motility and survival.

With the targeted *in vivo* CRISPR screening method developed here, we assessed the fitness contributions of 22 *Cryptosporidium* genes. This number is near to the total number of *Cryptosporidium* genes with a genetically verified impact on parasite virulence or survival, prior to this study. As this screening technology develops and improves, we anticipate greater throughput, allowing for the assessment of a variety of phenotypes on a larger scale. This rapid assessment of phenotypes will allow us to expand the knowledge base for *Cryptosporidium* and explore basic biology that is essential for developing more effective interventions.

## Methods

### Plasmid design and construction

Genomic sites for CRISPR directed repair were predicted using a customised version of EuPaGDT that retrieved 50bp of flanking DNA surrounding the predicted PAM site (Alvarez-Jarreta et al., 2023). Golden Gate Assembly or Gibson Assembly was used to generate all vectors used in this work. Golden Gate Assembly used, BsaI, BbsI-HF or BsmBI (New England BioLabs (NEB)) restriction enzymes. Gibson Assembly used HiFi DNA Assembly (NEB). To generate KO vectors for CRISPR screening, 2 consecutive Golden Gate reactions were performed. Firstly, between a Cas9-U6 vector and a 300bp unique segment containing 50bp homology arms (one of which contained the 50bp gRNA), tracrRNA and a DNA barcode, creating a Cas9-unique plasmid. Secondly, between this Cas9-unique plasmid and an interchangeable selection cassette, in turn creating the KO vector. To generate diCRE mediated inducible knockouts, a gene of interest (GOI) was recodonised and cloned into a LoxP-Nluc-NeoR-diCRE vector via Gibson Assembly. To generate Cas9-U6-gRNA vectors, a 20bp gRNA was cloned into the Cas9-U6-BsaI vector via Golden Gate Assembly (Vinayak et al., 2015), (Pawlowic et al., 2017).

### Culturing host cells

Human ileocecal adenocarcinoma cells (HCT8) were cultured in RMPI-1640 medium (Gibco) supplemented with 10% heat-inactivated foetal bovine serum (Merck), 120U/mL penicillin (Life Technologies) and 0.1% amphotericin B (Gibco) at 37°C under 5% CO_2_. Cells were passaged at approximately 70% confluence using 0.25% trypsin-EDTA (Gibco). Cells were used for experiments between passage numbers 5 and 25. For super resolution and ultrastructure expansion microscopy, cells were seeded in 24-well plates containing coverslips. For high throughput microscopy quantifications, cells were seeded in black 96-well clear bottom tissue culture-treated plates (Corning). For transient transfections using luminescence, cells were seeded in 24-well tissue culture-treated plates (Corning). For live imaging, cells were seeded in μ-slide 8-well high chambers (Ibidi).

### Oocysts and excystation

*C. parvum* IOWAII strain oocysts were purchased from Bunch Grass Farm (Deary, ID). Oocysts were stored at 4°C and used within 3 months of the date of isolation. The oocysts were excysted by incubating on ice with 1% sodium hypochlorite (VWR) in H_2_0 for 5 minutes, followed by incubating with 0.75% sodium taurocholate (Merck) in RPMI-1640 medium with 1% foetal bovine serum at 37°C (10 minutes for cell monolayer infections without transfection and 50 minutes prior to transfection). For *in vitro* infections, primed oocysts were used to infect HCT8 monolayers. To proceed with transfections, more than 60% of oocysts had to have excysted, observed using light microscopy at 50 minutes post sodium taurocholate treatment (Eclipse TS2R Nikon). For Supplementary Figure 1, 0.75% sodium taurocholate was substituted with either 0.75% sodium deoxycholate (Sigma) or 0.75% sodium taurodeoxycholate (Sigma), and 1% RPMI-1640 was substituted with PBS.

### Generation of transgenic parasites

Excysted sporozoites were suspended in Lonza SF buffer, combined with the appropriate DNA and electroporated using program ‘FL115’ on an AMAXA Nucleofactor 4D electroporator (Lonza) (FL115 was used for all transfections unless otherwise stated). For transient transfections *in vitro*, 5.0 x 10^6^ oocysts were excysted and sporozoites were transfected in the 20μL 16-well Nucleocuvette Strip format with 20μg of plasmid. For generating stable transgenic parasites *in vivo*, 2.5 x 10^7^ oocysts were excysted and sporozoites were transfected in the 100μL Nucleocuvette Vessel format with 20μg of Cas9-U6-gRNA plasmid and 40μg of repair cassette containing the 50bp homology arms. For CRISPR screens, 5.0 x 10^6^ sporozoites were excysted and sporozoites were transfected with 30μg of KO vector for each gene (when 2 KO vectors were used, 15μg of each KO vector was used). Transfections to knockout individual genes were performed separately (when 2 KO vectors per gene were used, transfections of KO vectors for the same gene were performed together). Parasites were pooled post transfection in 1% RMPI-1640 (Gibco) to infect mice and 4 or 5 mice were infected for each screen.

### Mouse model of infection

Interferon gamma deficient (*Ifnγ^-/-^*) mice were bred and housed in pathogen-free conditions in the Biological Research Facility at The Francis Crick Institute. Mice of both sexes were used for experiments. To increase infection efficiency, mice were pretreated with an antibiotic cocktail: 1g/L ampicillin (Merck), 1g/L streptomycin (Merck) and 0.5g/L vancomycin (Cambridge Biosciences) in their drinking water for 3 to 10 days prior to infection with transfected sporozoites. Before infection, mice received saturated sodium bicarbonate (Thermo Fisher Scientific) via oral gavage to neutralise their stomach acid. A second oral gavage was then undertaken 5 minutes thereafter with transfected sporozoites. Neither oral gavage exceeded the maximum volume of 0.1mL per 10g of body weight. For selection of transgenic parasites, 16mg/mL paromomycin (BioServ) was administered in the mice’s drinking water. To induce excision when diCRE-expressing transgenic parasites were used *in vivo*, 0.05 mg/mL rapamycin (Stratech Scientific) was administered in the mice’s drinking water. No drug treatment lasted more than 3 consecutive weeks. During murine infections, faecal material was collected daily and stored at 4°C. All experiments involving mice was done under the care and supervision of the Francis Crick Institute veterinary and Biological Research Facility staff, under protocols approved under project license PP8575470.

### Measuring parasite shedding by nanoluciferase

Faecal material was collected and 20mg was lysed in faecal lysis buffer (50mM Tris-hydrochloric acid, 10% glycerol, 1% Triton X-100, 2mM dithiothreitol, 2mM ethylenediaminetetraacetic acid (EDTA)). The lysate was clarified and combined with an equal volume of a 1:50 Nano-Glo Luciferase Assay Substrate: Nano-Glo Luciferase Assay Buffer (Promega). Luminescence was read at 200 gain on the BioTek Cytation5 (Agilent Technologies).

### Isolation of oocysts from mouse faeces

Faecal collections from the peak of infection were pooled, combined with cold water and filtered through a 250mm mesh. The faecal suspension was then mixed 1:1 with saturated sucrose and pelleted by centrifugation at 1000g for 10 minutes. Oocysts, located in the supernatant, were washed with cold water and pelleted by centrifugation at 1000g for 5 minutes. A caesium gradient was used to isolate 750μL of oocysts. Pure oocysts were washed with saline and stored for up to 6 months at 4°C in saline.

### CRISPR screening barcoding

To obtain barcodes from the output, barcodes could either be extracted from the peak of infection or directly from daily faecal samples. To do so from the peak of infection, oocysts were purified from pooled faecal material (5 to 7 days surrounding the peak of infection), excysted and gDNA was extracted using the DNeasy Blood & Tissue Kit (Qiagen). To do so from daily faecal samples, DNA was extracted from 100mg of faeces using the QIAamp PowerFecal Pro DNA Kit (Qiagen). Barcodes were obtained from the input material (100μL of the pool of transfected parasites used to infect the mice) using the QIAquick PCR Purification Kit (Qiagen). Barcodes from the input and output were amplified with high fidelity KAPA polymerase (Roche) (25 cycles, annealing 68°C and extension 15 seconds), and amplicon sequencing flanks were added with high fidelity KAPA polymerase (Roche) (10 cycles, 68°C annealing and 15 seconds extension). The success of the nested PCR was confirmed via gel electrophoresis and barcodes were submitted for amplicon sequencing to The Genomics Science Technology Platform at The Francis Crick Institute. Illumina MiSeq platform with a paired-end 250bp run configuration on a Nano flow cell was used.

### CRISPR screening analysis

The 7bp barcode between the barcode primer binding sites was bioinformatically extracted from both input and output samples. Barcodes for each gene were counted and the percentage of the barcode in the pool calculated for both the input and output. Using the % barcode in the pool for the input and output, fold changes of all genes were calculated:

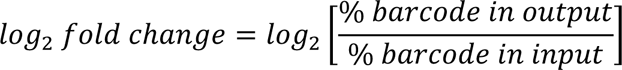

### Immunofluorescence microscopy

At the required timepoint, HCT8 cell monolayers were washed with 1X PBS, fixed with 4% PFA/PBS (Alfa Aesar) for 15 minutes, permeabilised with 0.25% Triton X-100 (Merck) for 10 minutes and blocked with 4% BSA/PBS (Merck) overnight at 4°C. Primary antibodies were incubated in 1% BSA/PBS for 2 hours and then cell monolayers washed 5 times with 1X PBS. Secondary antibodies were incubated for 1 hour in 1% BSA/PBS along with the fluorescein labelled 1:4000 Vicia villosa lectin (VVL) (Vector Lab) or 1:1000 Helix pomatia agglutinin (HPA) (Invitrogen) that stains parasites. Nuclei were stained by incubating with 1:10,000 Hoechst 33342 (Invitrogen) in 1X PBS for 5 minutes. For the EdU assays, 10mM EdU was incubated from 28 to 32 hours post infection and stained as the described in the Click-iT EdU Cell Proliferation Kit for Imaging (C10340, Invitrogen). Stained monolayers were washed with 1X PBS, the coverslips mounted with ProLong Gold Antifade (Thermo Fisher Scientific) and visualised using a VisiTech instant super resolution imaging system (VT-iSIM). Alternatively, when super resolution was not required, the BioTek Cytation5 (Agilent Technologies) was used to visualise the stained monolayer. See **Table 6** for a full list of antibodies used in this work.

**Table 3.**
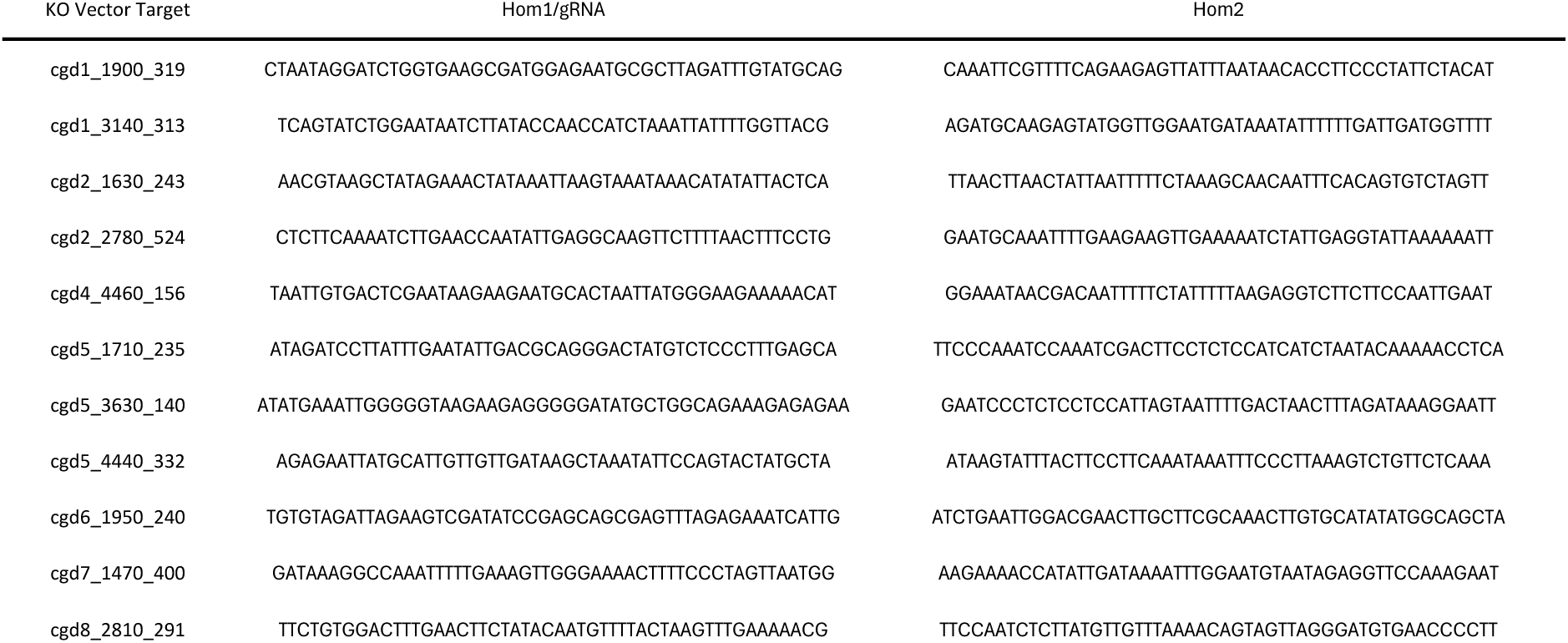
Homology Arms/gRNAs used in Pyrimidine Salvage 1 KO Vector CRISPR Screen.

**Table 4.**
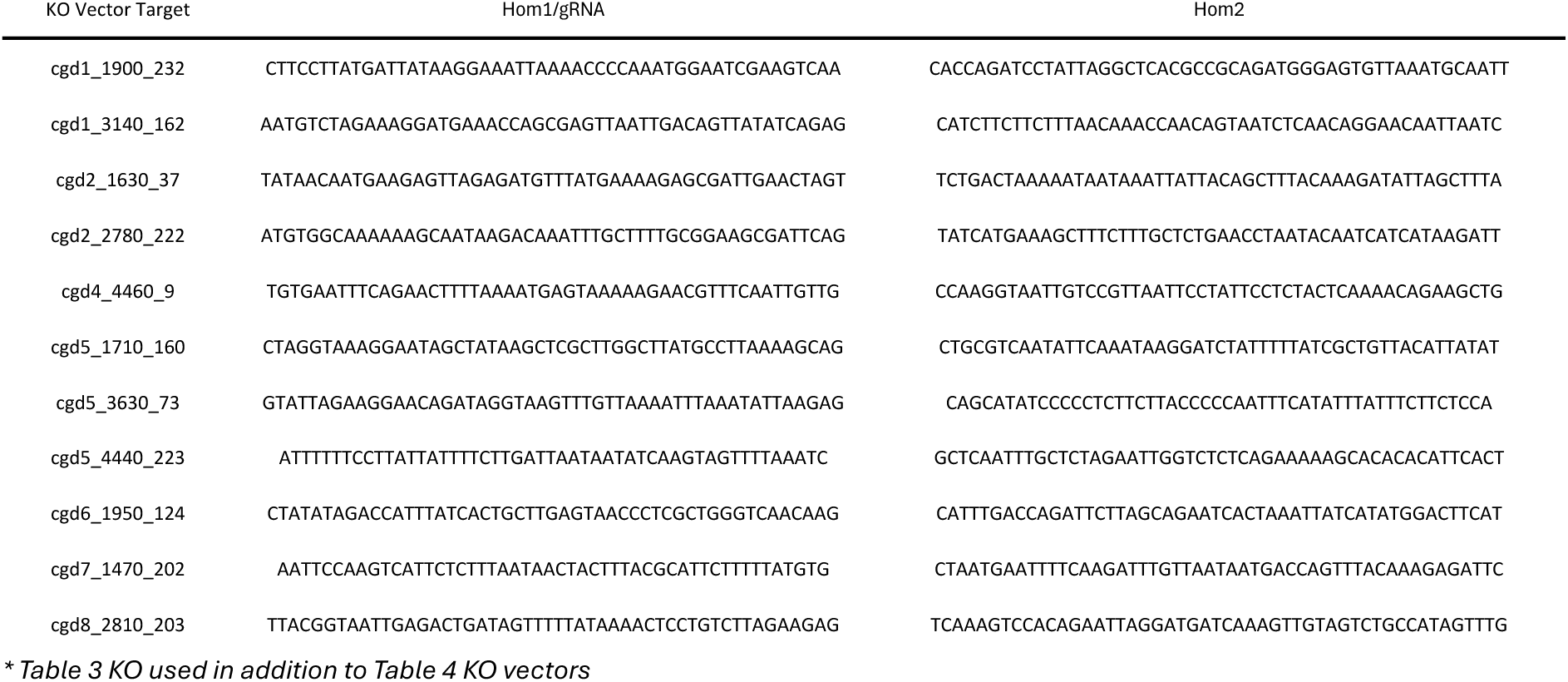
Homology Arms/gRNAs used in Pyrimidine Salvage 2 KO Vector CRISPR Screen.

**Table 5.**
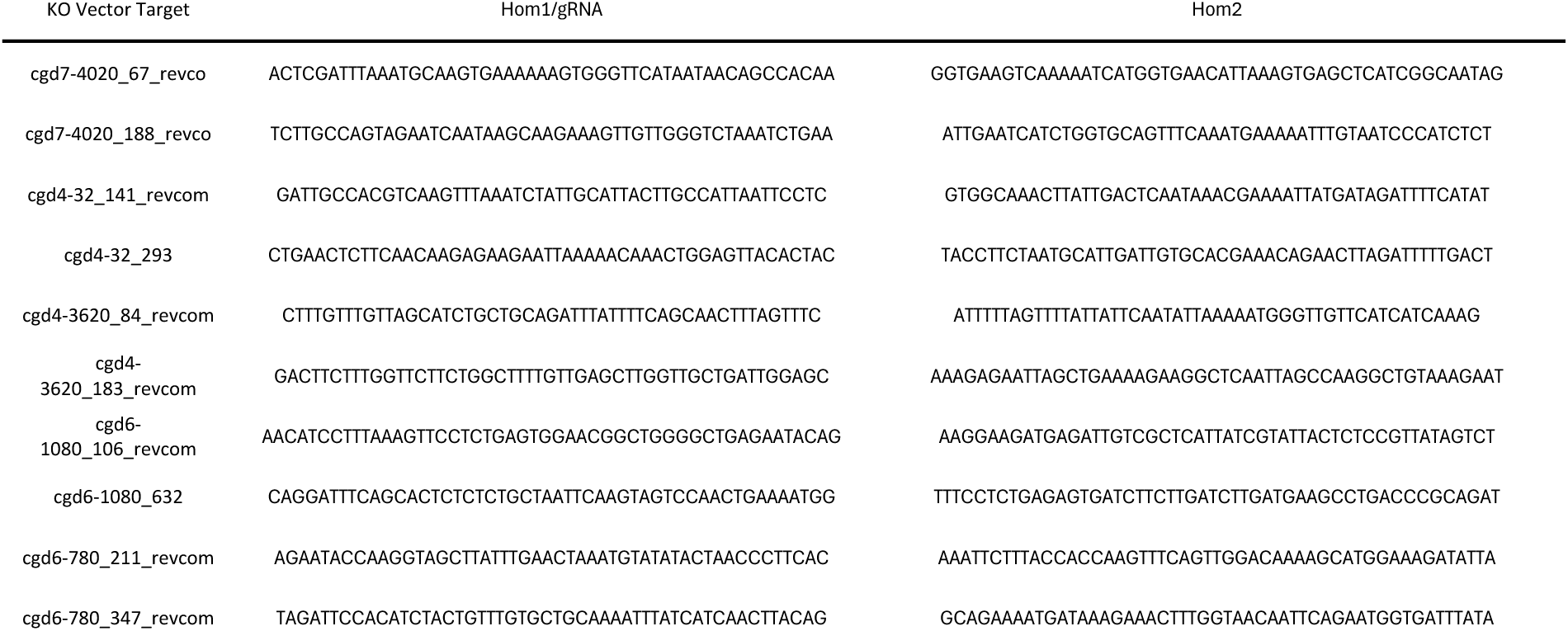
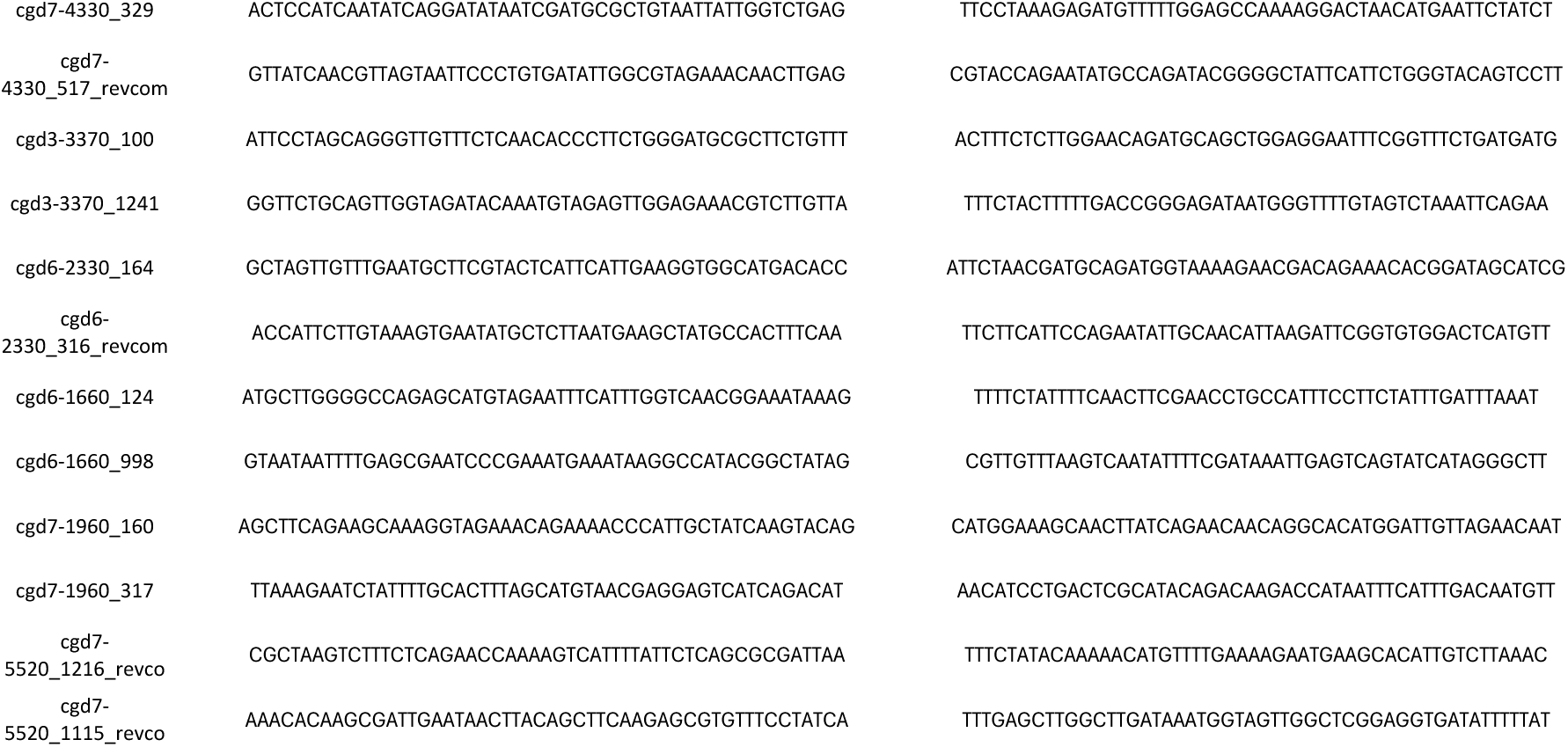
Homology Arms/gRNAs used in Vaccine Candidate 2 KO Vector CRISPR Screen.

**Table 6.**
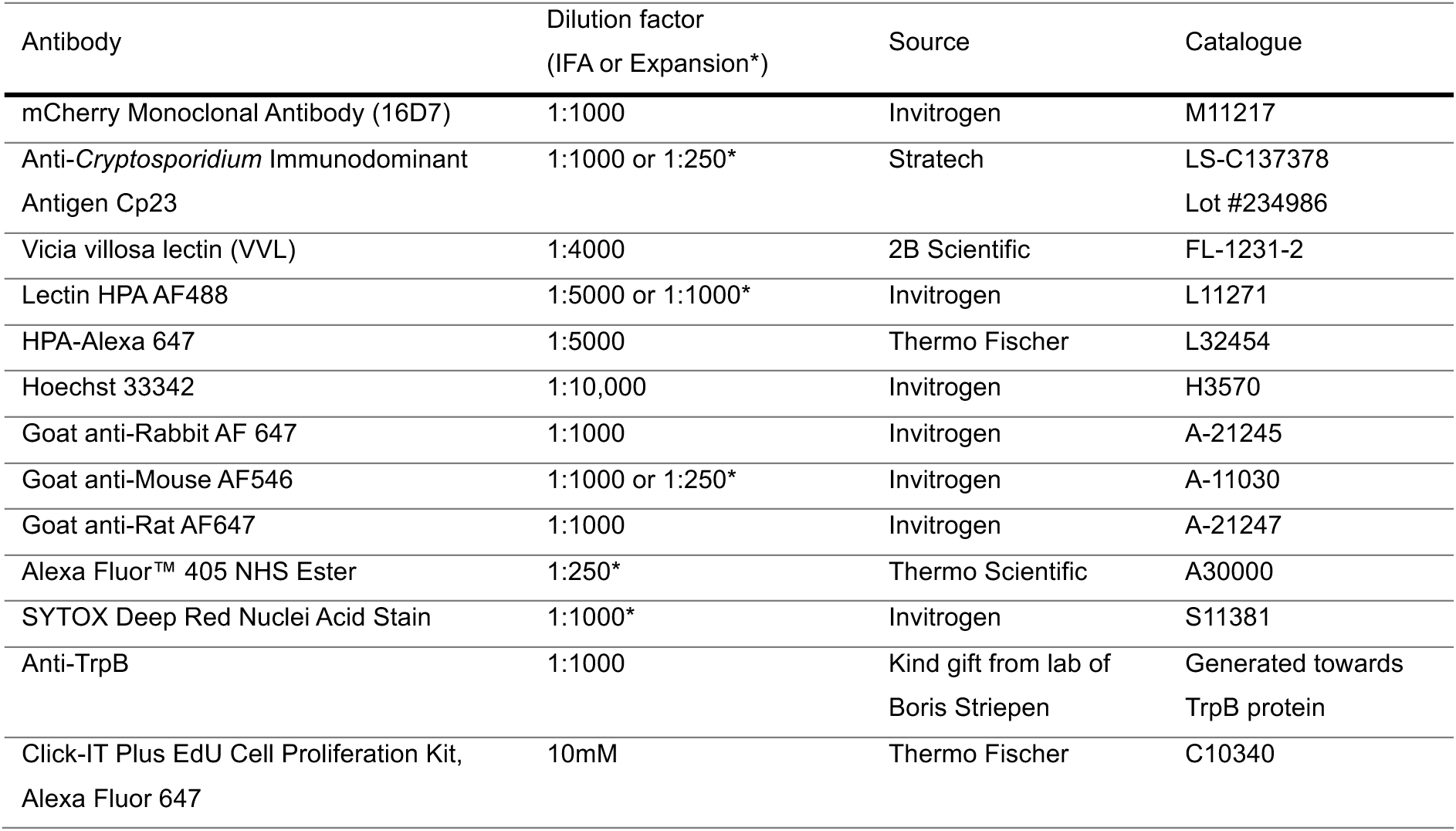
List of Antibodies Used in this Work.

### Ultrastructure expansion microscopy

This protocol has been adapted from Liffner and Absalon, 2021. For the expansion of sporozoites, coverslips were coated with 0.1mg/mL poly-D-lysine for 1 hour and then washed twice with 1X PBS. Excysted sporozoites in 1% RPMI-1640 were added and allowed to adhere for 10 minutes at 37°C. Adhered sporozoites or cell monolayers to expand were fixed with 4% PFA/PBS for 15 minutes at 37°C. Protein crosslinking prevention was performed by adding 1.4% formaldehyde/ 2% acrylamide in 1X PBS to each sample, then incubating overnight at 37°C. To perform gelation, TEMED and APS were added to a monomer solution (19% w/w sodium acrylate / 10% v/v acrylamide / 0.1% v/v N,N’-methylenebisacrylamide in 1X PBS) and pipetted under each coverslip in a pre-cooled humid chamber which was then incubated for 5 minutes on ice. To complete gelation, the humid chamber was incubated for 1 hour at 37°C. To separate gels from the coverslips, the coverslips were incubated with a denaturation buffer (200mM SDS, 200mM NaCl, 50mM Tris in water, pH 9) for 15 minutes. To complete the denaturation of the sample, the gel was incubated in a denaturation buffer for 90 minutes at 95°C. To expand the sample, the gel was incubated with dH_2_0 for 30 minutes, three times in total at room temperature (RT). Following the first round of expansion, the sample was shrunk by incubating with 1X PBS, 2 times in total at RT. To block the sample, 2% BSA/PBS was incubated for 1 hour at RT. To stain the sample, primary antibodies were incubated in 2% BSA/PBS overnight at RT. Gels were then washed 3 times in total with 0.5% Tween 20/PBS for 10 minutes. Directly conjugated and secondary antibodies were incubated in 1X PBS for 3 hours at RT. The gel was then washed 3 times in total with 0.5% Tween20/PBS for 10 minutes. A second round of expansion took place by incubating the gel with dH_2_0 for 30 minutes, three times at RT. The gel was measured to calculate the expansion factor, mounted onto a 0.1mg/mL poly-D-lysine coated 60mm dish and visualised on the VT-iSIM microscope.

### Immunogold electron microscopy

1x10^7^ oocysts were excysted for 1 hour, pelleted and re-suspended in fixative (8% formaldehyde in 0.4M HEPES buffer, pH 7.4) for 15 minutes at RT. Samples were washed with 0.2M HEPES and a secondary fixation (2% formaldehyde + 0.05% glutaraldehyde in 0.2M HEPES) step was carried out at 4°C overnight. Samples were dehydrated and infiltrated with LR white resin at -20°C overnight. Prior to polymerisation, samples were brought back to RT for 1 hour and then polymerised at 60°C for 24 hours. The samples were sectioned using a Leica UC7 ultramicrotome with a 45° Diatome diamond knife, achieving sections of 70nm thickness. The sections were collected on nickel grids and immunogold labelled. To do so, the samples were quenched with PBS/glycine for 2 minutes three times and then blocked with 1% BSA/PBS for 5 minutes at RT. A 1:10 dilution of the primary Cp23 antibody was incubated in 1% BSA/PBS for 1 hour and then samples were washed twice with 0.1% BSA/PBS. A 1:50 dilution of the protein-A gold bound to 10nm gold particles (PAG-10) was incubated with the sample in 0.1% BSA/PBS for 20 minutes and then washed twice with PBS. A post fixation step was carried out with 1% glutaraldehyde for 5 minutes and samples were washed with Milli-Q H_2_0 for 1 minute 6 times in total. Samples were incubated in 1% uranyl acetate for 10 minutes and air dried. Transmission electron microscopy was carried out on the 120-kV JEOL JEM-1400Flash Electron Microscope (JEOL Ltd., Welwyn Garden City, UK) with a JEOL Matataki Flash camera.

### Live microscopy

At 18 hours post-infection, phase contrast imaging was performed using an Eclipse Ts2R microscope (Nikon) with a 40X/0.55 NA Ph1 ADL objective (Nikon), digital sight 10 camera (Nikon) and a TPi-TCSX (Tokai Hit) heated stage set at 37°C. Images were acquired at 25fps for 1 hour to capture egressing merozoites. Images were imported into ImageJ2 Version 2 and the number of gliding merozoites was manually observed. Image sequences were converted to movies to acquire short videos of egressing merozoites.

### Permeabilisation assay

Coverslips placed in 24-well plates were coated with 0.1mg/mL poly-D-lysine for 1 hour and washed twice with 1X PBS. Excysted sporozoites in serum free RMPI-1640 were allowed to adhere for 10 minutes at 37°C. Adhered sporozoites were fixed with 1% PFA/PBS for 20 minutes at RT. For the permeabilised condition, 0.1% Triton X-100 was used to permeabilise for 10 minutes at RT. For the non-permeabilised condition, 1X PBS was instead incubated for 10 minutes at RT. Both conditions were blocked with 4% BSA/PBS overnight at 4°C. Primary antibodies were incubated in 1% BSA/PBS for 2 hours at RT. The coverslips were washed 5 times and secondary antibodies were incubated in 1% BSA/PBS for 1 hour at RT. All coverslips were mounted with ProLong Gold Antifade (Thermo Fisher Scientific) and visualised using a VT-iSIM.

### Attachment assay

Coverslips placed in 24-well plates were coated with 0.025mg/mL poly-D-lysine for 1 hour and washed twice with 1X PBS. 60,000 oocysts per well were excysted and sporozoites were added and spun at 80g for 1 minute to adhere. Attachment was performed in Ringer’s solution (10mM Hepes pH 6.7, 10mM Glucose, 2mM CaCl_2_, 1mM MgCl_2_, 3mM KCl, 3mM NaH_2_PO_4_ and 155mM NaCl) for 15 minutes at 37°C. Sporozoites were fixed using 8% PFA/PBS leak in resulting in 4% PFA/PBS/Ringer’s fixation for 15 minutes at RT. Sporozoites were washed 3 times in total with 1X PBS and blocked with 4% BSA/PBS overnight at 4°C. Sporozoites were stained with 1:5000 Helix pomatia agglutinin (HPA) in 1% BSA/PBS for 1 hour at RT. The BioTek Cytation5 (Agilent Technologies) was used to visualise the attached sporozoites. Images were exported as TIFs into ImageJ2 Version 2 and the below macro was used to analyse the data:

run(“Subtract Background…”, “rolling=50”);

setAutoThreshold(“Default dark”);

//run(“Threshold…”);

setAutoThreshold(“Triangle dark”);

setOption(“BlackBackground”, true);

run(“Convert to Mask”);

run(“Median…”, “radius=2”);

run(“Analyze Particles…”, “size=50-10000 circularity=0.00-1 display summarize overlay”);

### Neutralisation assay

HCT8 cells were seeded to confluency in 96-well clear bottom tissue culture-treated plates (Corning). An 8-point 2-fold dilution series of the Cp23 antibody (Stratech, LS-C137378) and the IgG isotype control (Stratech, GTX35009 + 0.02% Proclin 300 (Merck, 48912-U) was prepared (0.02mg/mL to 0.0002mg/mL) in 96-well non-tissue culture-treated plates. 25,000 oocysts per well were excysted and incubated with the antibody dilution series for 5 minutes at 37°C. Post antibody incubation, the sporozoites were added to the 96-well plate containing HCT8 cells in 1% RPMI-1640. At 24 hours post-infection, cell monolayers were washed with 1X PBS, fixed with 4% PFA/PBS (Alfa Aesar) for 15 minutes, permeabilised with 0.25% Triton X-100 (Sigma) for 10 minutes and blocked with 4% BSA/PBS (Merck) overnight at 4oC. Parasites were stained with 1:4000 Vicia villosa lectin (VVL) and host nuclei were stained with 1:10,000 Hoechst 33342 (Invitrogen) in 1% BSA/PBS for 1 hour at RT. Cell monolayers were washed and visualised on the BioTek Cytation5 (Agilent Technologies). 2x2 tiled images per well were acquired with the 20X objective. The Gen5 analysis software (Agilent Technologies) was used to count the number of host cell nuclei and parasites for parasite per nuclei calculations.

## Supporting information

Video_S1_DMSO

Video_S2_DMSO

Video_S3_RAP

Video_S4_RAP

## Acknowledgements

We would like to thank Pippa Hawes of The Francis Crick Electron Microscopy Scientific Technology Platform for her helpful discussions and Elena Rodrigues of the Cryptosporidiosis Laboratory for preparing *Cryptosporidium* samples for electron microscopy. Further, we would like to thank Nicholas Chisholm and other members of the Biological Research Facility, as well as the Genomics Scientific Technology Platform at The Francis Crick Institute for their contributions to this work. We would also like to thank Rodrigo Baptista of Houston Methodist for his insight and helpful conversations. This work was supported by the Francis Crick Institute—which receives its core funding from Cancer Research UK, the UK Medical Research Council, and the Wellcome Trust (CC2063) – and a UKRI grant awarded to A.S. (101042783).

## Author Contributions

L.C.W & A.S, with help from K.S, N.B, L.B & N.B.M, pioneered the CRISPR screening approach. M.P developed the attachment assay. L.C.S & L.C carried out TEM. D.P developed the customised version of EuPaGDT used for predicting CRISPR guides. L.C.W and A.S wrote the manuscript with contribution from all of the authors.

---

**Supplementary 1.**
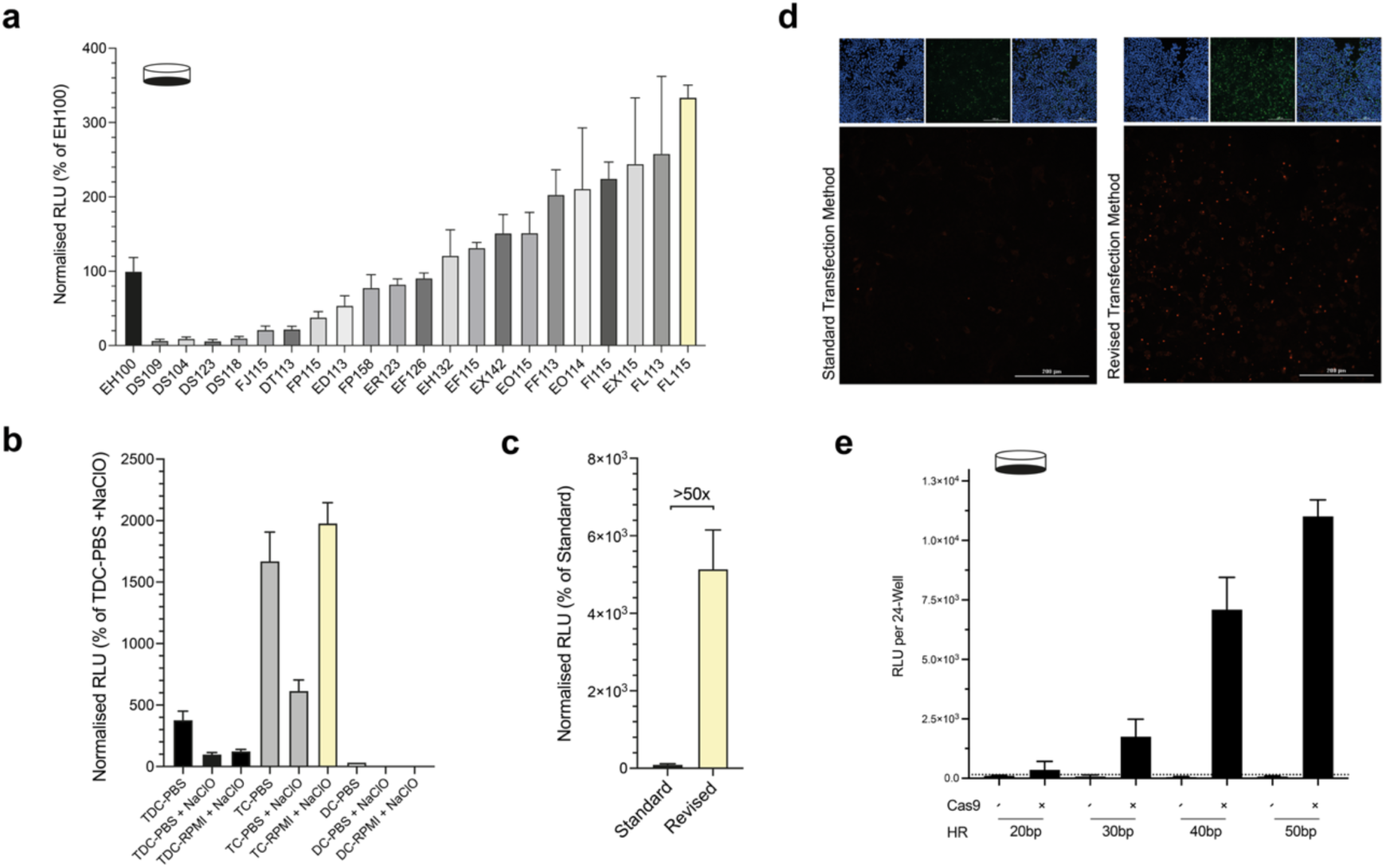
Advancement to CRISPR Screening. **a-b.** Relative luminescence units (RLU) from a HCT8 cell monolayer infected for 24 hours with *C. parvum* parasites that were transiently transfected with a nanoluciferase-mCherry expression vector, trialling multiple electroporation programs (**a**) and excystation reagents used prior to transfection (**b**). TDC (taurodeoxycholate), NaCIO (sodium hypochlorite), TC (taurocholate), RMPI (media), DC (deoxycholate) PBS (phosphate-buffered saline). **c**. Quantification of the improved transfection efficiency measured by RLU. In black, the standard method, in yellow, the revised method. **d.** Visualisation of the increased transfection efficiency of the revised method (right panel) compared to the standard method (left panel) using the mCherry-expression vector. Blue (Hoechst), nuclei; green (Vicia villosa lectin (VVL)), parasites; red (mCherry Ab), mCherry/transfected parasites. All data shows the mean ± standard deviation from 2 biological replicates. **e.** RLU from a HCT8 cell monolayer infected for 24 hours with *C. parvum* parasites transfected with a repair cassette designed to replace the thymidine kinase gene, using different lengths of homology repair. Data shows the mean ± sd from three technical replicates.

**Supplementary 2.**
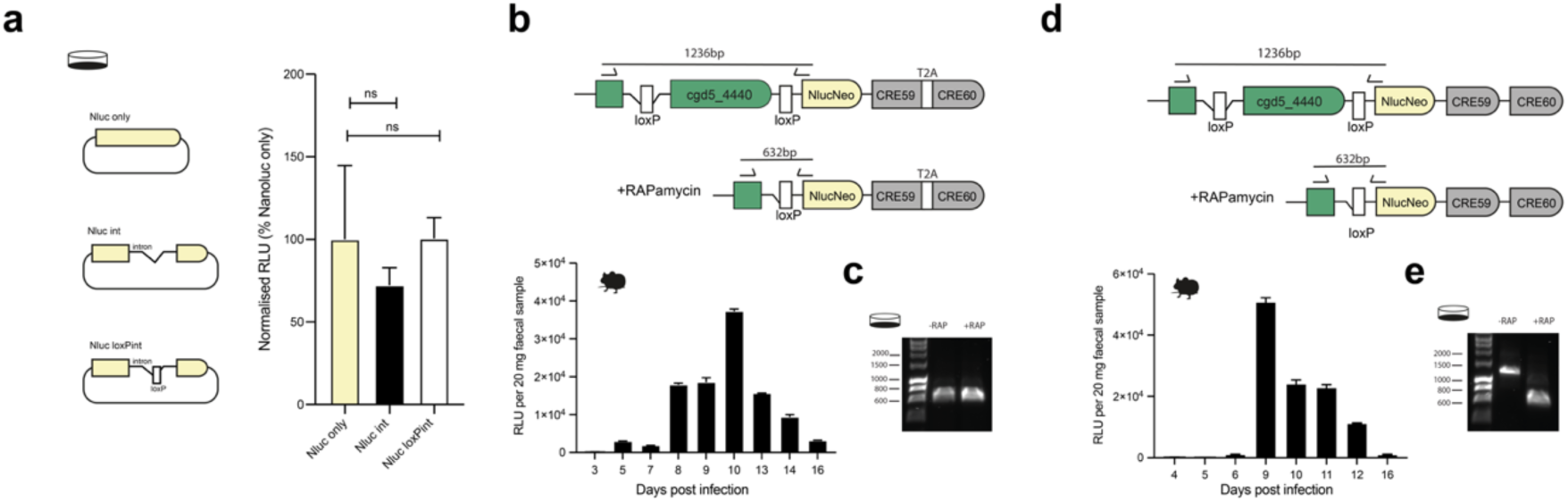
Refining diCRE Mediated Excision. **a.** Schematics of disruption of a nanoluciferase (Nluc) gene mid-codon with a HAP2 (cgd8_2220) intron (Nluc int) or with a HAP2 intron containing a loxP sequence (Nluc loxPint). Vectors were transiently transfected into *C. parvum* sporozoites that were allowed to infect an HCT8 monolayer for 24 hours. Data shown is the mean of 2 biological replicates ± sd of 4 technical replicates. Significance is determined using a one-way ANOVA, ns = not significant **b.** Schematic of the TK-T2A-diCRE parasite line and the corresponding luminescence from mouse faecal material when generating these parasites. Data shown is a biological replicate ± SEM, n = 4 Ifnγ^-/-^ mice. **c.** Diagnostic PCR to determine the level of excision occurring in the TK-T2A-diCRE parasite line after 24 hours infection of a HCT8 cell monolayer during treatment with rapamycin. **d.** Schematic of the TK-diCRE parasite line and the corresponding luminescence from mouse faecal material when generating the parasites. Data shown is a biological replicate ± SEM, n = 4 Ifnγ^-/-^ mice. **e.** Diagnostic PCR to determine the level of excision occurring in the TK-diCRE parasite line after 24 hours infection of a HCT8 cell monolayer during treatment with rapamycin.

**Supplementary 3.**
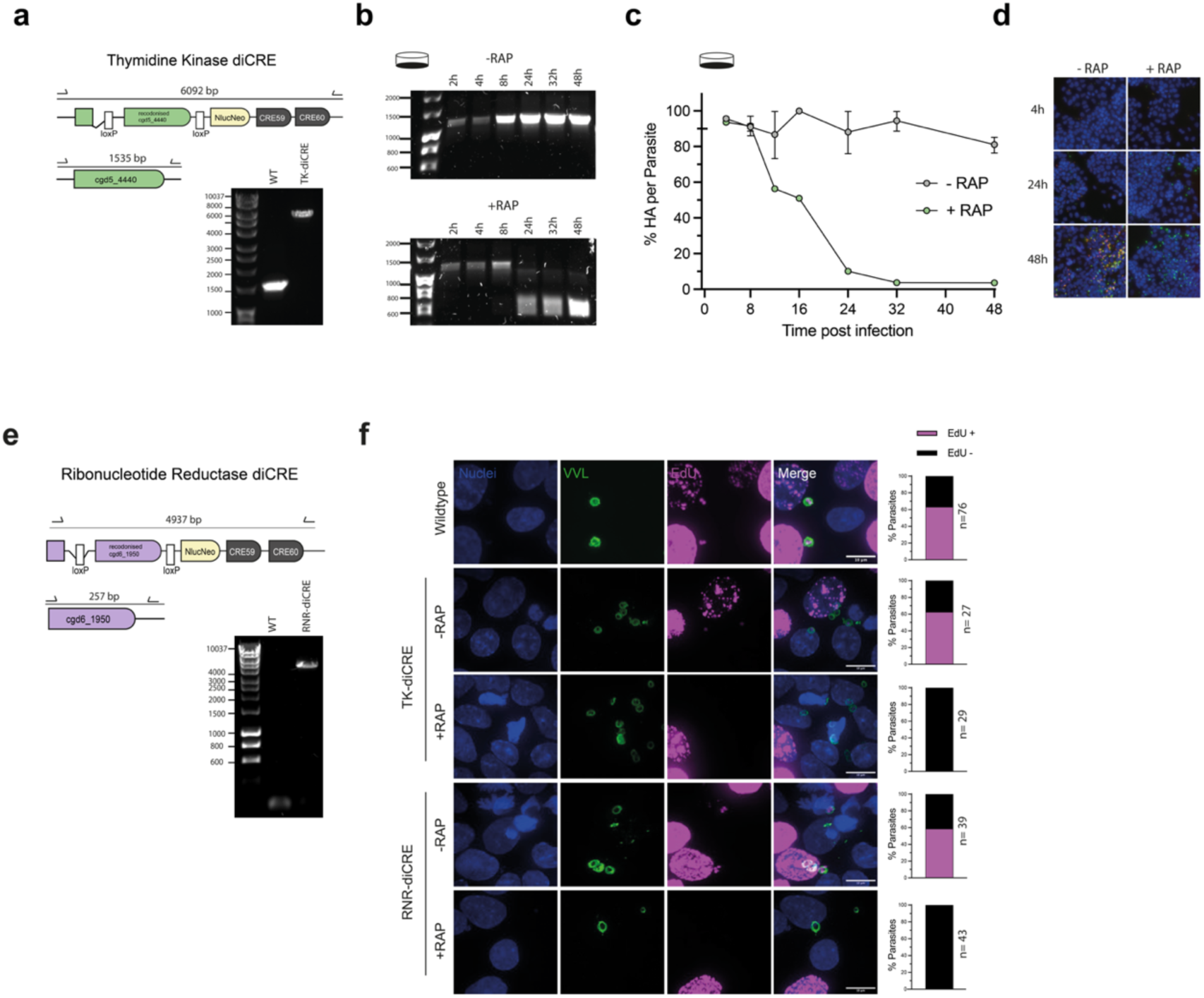
diCRE Mediated Excision of TK and RNR. **a.** Schematic and genotyping PCR of TK-diCRE parasites. **b.** TK-diCRE parasites were allowed to infect a HCT8 monolayer and genomic DNA was extracted at 2, 4, 8, 24, 32 and 48 hours post infection in the presence and absence of rapamycin. Diagnostic PCRs confirmed the level of excision that had occurred at the respective timepoint. **c-d.** TK-diCRE parasites were allowed to infect a HCT8 monolayer in the presence and absence of rapamycin, and the monolayer was fixed and stained at 4, 8, 12, 16, 24, 32 and 48 hours post infection. The HA tag per parasite was quantified by automated imaging, representation images are shown in **d**; blue (Hoechst); green (Vicia villosa lectin (VVL)); red (αHA). Data shown is representative of 2 biological replicates, each with 2 technical replicates ± sd. **e.** Schematic and genotyping PCR of RNR-diCRE parasites. **f.** EdU assay with quantifications showing newly synthesised DNA between 28 and 32 hours in wildtype, TK- and RNR-diCRE parasites in the presence and absence of rapamycin. Blue (Hoechst); green, (Vicia villosa lectin (VVL)); magenta, (EdU). Data shown is two biological replicates with the number of parasites quantified indicated. Scale bar = 10μm.

**Supplementary 4.**
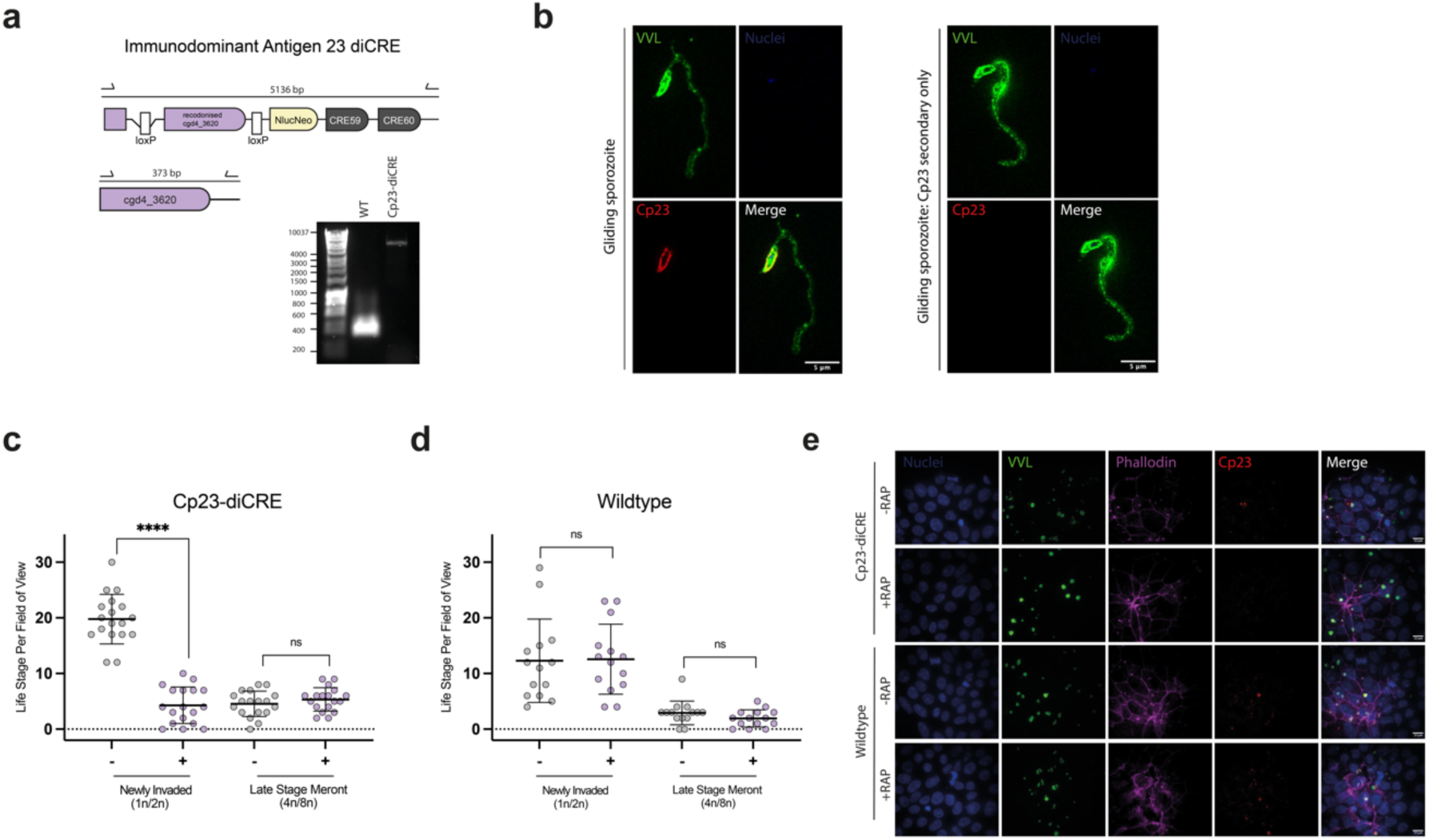
Immunodominant Antigen 23 is Essential and Required for Reinvasion of Host Cells. **a.** Schematic and genotyping PCR of Cp23-diCRE parasites. **b.** Visualising gliding *C. parvum* sporozoites using super-resolution microscopy. Green (Vicia villosa lectin (VVL) which marks both parasite and trail); red (αCp23); blue (Hoechst). Scale bar = 5 μm. **c-e.** Cp23-diCRE and wildtype parasites were allowed to infect an HCT8 monolayer in the presence or absence of rapamycin. At 22 hours post infection the life cycle stage was quantified. Data shows 2 biological replicates. Significance is determined using a two-tailed unpaired t-test, ns = not significant, **** = p ≤ 0.0001. Representative images are shown in **e**. Blue (Hoechst); green (Vicia villosa lectin (VVL)); magenta (phallodin - actin); red (αCp23). Scale bar = 10μm.

**Supplementary 5.**
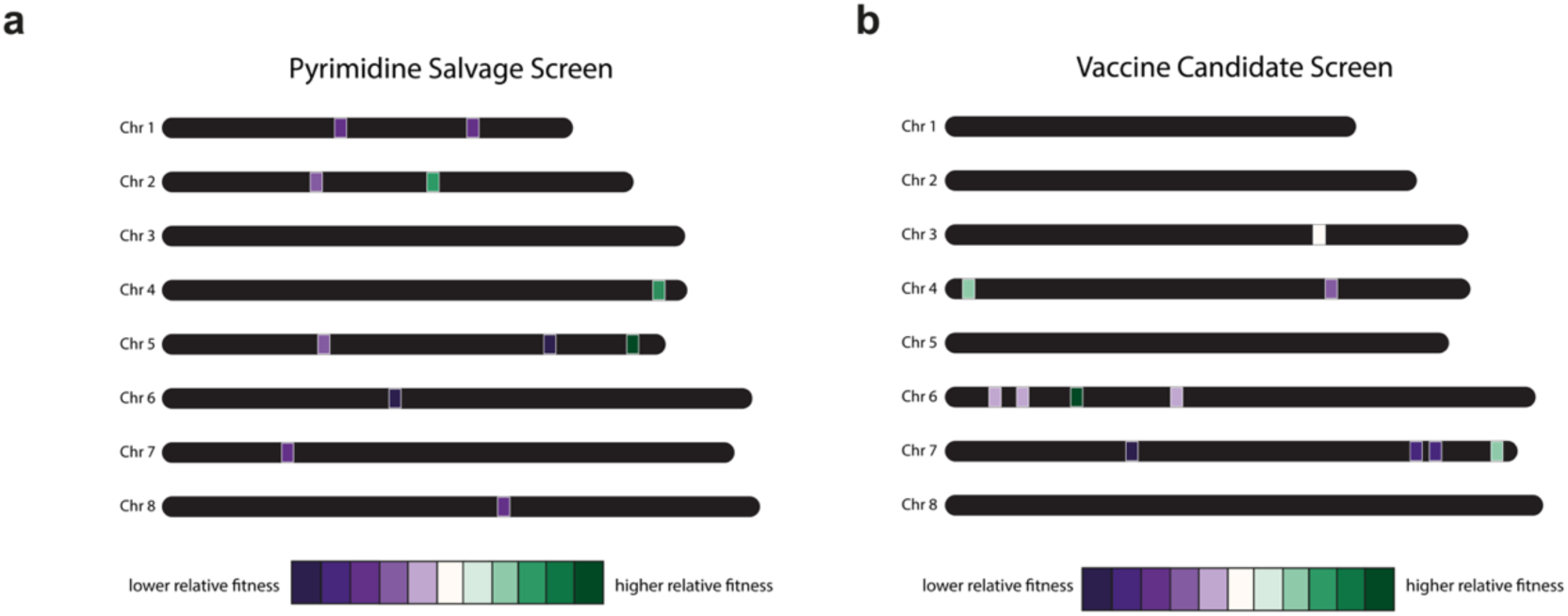
Chromosome Locations of the Genes Knocked Out During CRISPR Screens. **a-b.** Locations of the genes from the pyrimidine salvage (**a**) and vaccine candidate (**b**) CRISPR screens. The colour indicates the relative fitness of the gene knocked out with dark purple being a highly fitness conferring gene and dark green being a low fitness conferring gene.

**Supplementary 6.**
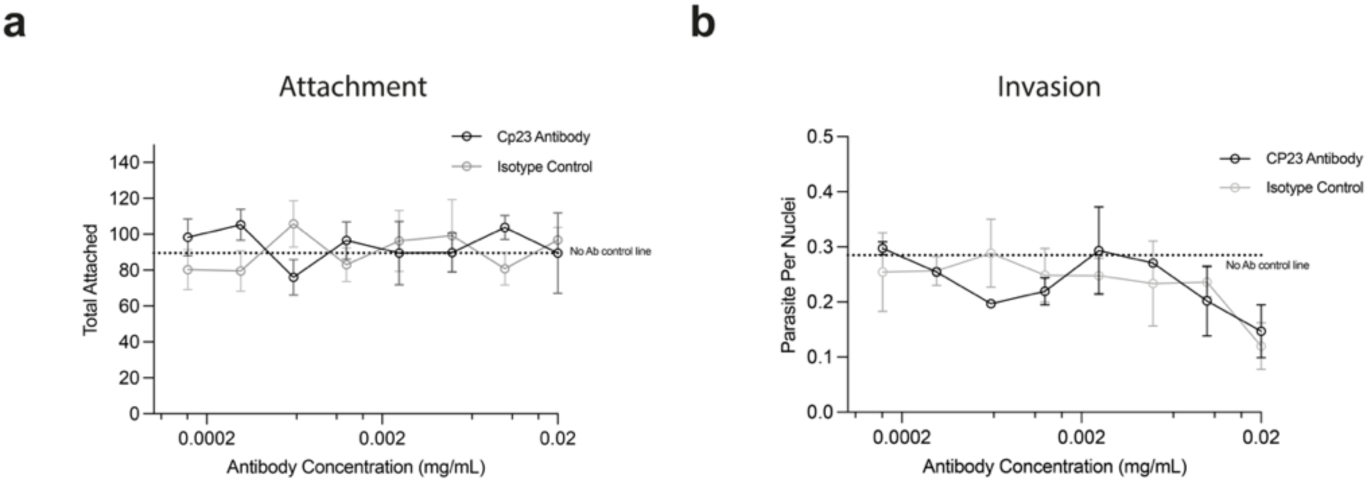
Immunodominant Antigen 23 Antibody Does Not Neutralise Infection *In Vitro*. **a.** Quantifying the number of attached sporozoites on a poly-D treated surface when incubated with the Cp23 antibody or isotype control antibody dilution series. Data shows the mean ± sd of 4 technical replicates. The black line shows Cp23 antibody incubations, the grey line shows isotype control antibody incubations, and the dotted line shows no antibody control incubations. **b.** Quantification of invasion (parasite per host nuclei) when cell monolayers are infected in the presence of the Cp23 antibody or isotype control. The data shows the mean and ± sd of the 4 technical replicates. The back line shows Cp23 antibody incubations, the grey line shows isotype control antibody incubations, and the dotted line shows no antibody control incubations.

**Supplementary videos.** Live imaging of the reinvasion event was carried out at 18 hours post infection of an HCT8 monolayer with Cp23-diCRE parasites in the presence of rapamycin or vehicle (DMSO) control.

**Video S1. DMSO treatment 1**

**Video S2. DMSO treatment 2**

**Video S3. Rapamycin treatment 1**

**Video S4. Rapamycin treatment 2**

## Notes

### Competing Interest Statement

The authors have declared no competing interest.

